# A theory of oligogenic adaptation of a quantitative trait

**DOI:** 10.1101/2023.04.20.537719

**Authors:** Ilse Höllinger, Benjamin Wölfl, Joachim Hermisson

## Abstract

Rapid phenotypic adaptation is widespread in nature, but the underlying genetic dynamics remain controversial. Whereas population genetics envisages sequential beneficial substitutions, quantitative genetics assumes a collective response through subtle shifts in allele frequencies. This dichotomy of a monogenic and a highly polygenic view of adaptation raises the question of a middle ground, as well as the factors controlling the transition. Here, we consider an additive quantitative trait with equal locus effects under Gaussian stabilizing selection that adapts to a new trait optimum after an environmental change. We present an analytical framework based on Yule branching processes to describe how phenotypic adaptation is achieved by collective changes in allele frequencies at the underlying loci. In particular, we derive an approximation for the joint allele-frequency distribution at threshold levels of the trait mean as a comprehensive descriptor of the adaptive architecture. Depending on the model parameters, this architecture reproduces the well-known patterns of sequential, monogenic sweeps, or of subtle, polygenic frequency shifts. Between these endpoints, we observe oligogenic architecture types that exhibit characteristic patterns of partial sweeps. We find that a single compound parameter, the population-scaled background mutation rate Θ_bg_, is the most important predictor of the type of adaptation, while selection strength, the number of loci in the genetic basis, and linkage only play a minor role.

## Introduction

Quantitative traits (QTs) are everywhere in nature, from body size and milk yield to melanism and life expectancy. This has made the study of their evolution a subject of interest for more than a century (Walsh and Lynch 2018; Sella and Barton 2019). Ever since the modern synthesis, it has been clear that even complex QTs are governed by a finite genetic basis (Fisher 1918). Nevertheless, the genomic changes underlying phenotypic adaptation have long been elusive. Instead, much effort has gone into theoretical exploration. Two schools of thought – population genetics and quantitative genetics – developed independent narratives for the genetics of adaptation, with notable differences.

On the one hand, population genetics is concerned with the dynamics of allele frequencies. Consequently, adaptation is understood as major changes in allele frequencies driven by selection. The archetypal scenario is a (hard) selective sweep, where an initially rare beneficial allele at a single locus quickly rises to fixation (Maynard-Smith and Haigh 1974; Kaplan *et al*. 1989; Barton 1998). More complex models examine sweep patterns in diversity and divergence data across multiple loci in the basis of a trait or along a biological pathway (*e*.*g*., Gouy *et al*. 2017), discuss linkage and epistasis, and the effect of recurrent sweeps on genome-wide diversity levels observable through genome scans (reviewed in Stephan 2019). At the level of phenotype or fitness, this results in a view of adaptation via successive selective sweeps as discrete steps of a so-called adaptive walk (reviewed in Orr 2005).

On the other hand, quantitative genetics focuses on phenotype data. It thrives on being able to abstract from the underlying genetics, which only enter as summary statistics across loci, such as the genetic variance and higher moments of the trait distribution (Turelli and Barton 1990; Bürger 2000). Accordingly, the dynamics at single loci do not play a role. This view is taken to the extreme in the infinitesimal model, where phenotypic adaptation in completely decoupled from the changes in underlying allele frequencies (Fisher 1918; Barton *et al*. 2017). At the genotypic level, adaptation is viewed as tiny allele frequency shifts at a myriad of loci with small individual effects, but these dynamics are not explicitly described.

After decades of research with little exchange between communities, the emergence of data from genome-wide association studies (GWAS) led to a growing interest in an integrated view linking phenotypic adaptation with its genetic underpinnings. Pritchard and colleagues (Pritchard *et al*. 2010; Pritchard and Di Rienzo 2010) proposed the concept of “polygenic adaptation” as an alternative to the selective sweep model with the goal of characterizing – and, if possible, detecting – patterns of QT adaptation at the genomic level.

If adaptation can be either highly polygenic or involve only single loci, this raises the question of a middle ground, *i*.*e*., whether there is room for an “oligogenic view of adaptation” (Bell 2009), with its own characteristic patterns. To address this question, a framework is needed that connects the opposite endpoints and also covers the parameter range in between. Several recent concept papers discuss such frameworks (Láruson *et al*. 2020; Barghi *et al*. 2020; Fagny and Austerlitz 2021).

It turns out that the decisive parameter for determining the type of adaptation is not simply the number of loci that underlies a trait. What matters is a measure of *redundancy* (Láruson *et al*. 2020; Barghi *et al*. 2020), which determines how many alternative genotypes that solve the adaptive task can be readily generated in a population (and therefore coexist), either from standing genetic variation (SGV) or new mutation. To make these notions precise, a model is needed.

Here, we consider a classical additive trait under Gaussian stabilizing selection that adapts to a new trait optimum after an environmental change. This is a classic scenario that has been studied many times before (*e*.*g*., Lande 1976; Jain and Stephan 2017b; Stetter *et al*. 2018; Thornton 2019; Hayward and Sella 2022). We ask how adaptive progress of the trait mean toward the new optimum breaks down into contributions of alleles at the underlying loci. Specifically, we develop an analytical theory to approximate the joint distribution of these allelic contributions across replicates.

Our method builds on a two-step approach first used in Höllinger *et al*. (2019) to describe adaptation of a binary polygenic trait (such as pesticide resistance). In the first step, a multi-type branching process is used to infer the joint frequency distribution of co-segregating alleles, as long as these frequencies are small and strongly affected by genetic drift, but hardly interact (both before and after the environmental change). The second step describes how this distribution is transformed when the frequencies grow larger. While genetic drift can be ignored in this phase, epistatic interactions begin to play a role. This transformation is straightforward for a binary trait, but it can be extended to a much larger class of models, including the additive QT, in particular.

In this manuscript, we describe this method for the case of a trait with arbitrarily many unlinked additive loci of equal effect. We derive the joint frequency distribution of all alleles that contribute to phenotypic adaptation at arbitrary threshold values of the mean trait. The results show how the genotypic patterns change with an increasing phenotypic distance and provide a comprehensive classification of adaptive architectures, from single, consecutive sweeps to small polygenic shifts. In particular, we describe and discuss the characteristics of oligogenic architectures that lie between the better known monogenic or highly polygenic endpoints.

The remainder of the article is organized as follows: In the Model and Methods section, we first introduce the model and the assumptions of the simulation methods (individual-based and Wright-Fisher). Then, both steps of the analytical approximation are explained, while all detailed derivations can be found in a separate *Mathematical Appendix*. In the Results section, we compare the analytical predictions with simulation results and describe the most important patterns for both short- and long-distance adaptation. We also assess the effects of linkage between selected loci. The Discussion examines key notions and highlights the scope and limits of the model. The Appendix shows complementary figures. Scripts and data are deposited on *Dryad* (DOI: 10.5061/dryad.573n5tbc9, Höllinger *et al*. 2023).

## Model and Methods

### The model

We model a panmictic population of size *N*_e_ and follow the adaptation of an additive QT, *Z*, that is governed by *L* loci. Each locus *i* ∈{1, 2, …, *L}* is biallelic with alleles *a*_*i*_ and *A*_*i*_ and equal effect sizes set to 0 (for *a*_*i*_) and *γ >* 0 (for *A*_*i*_). We focus on haploid genetics in the main text for simplicity, and present extensions to diploids in the Appendix (Section Diploids with linkage). With *η*_*i*_ ∈ {0, 1} indicating the allelic state, we thus have

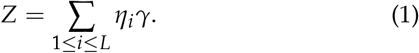

Wrightian fitness is modeled by time-dependent Gaussian stabilizing selection towards a trait optimum *Z*_opt_(*t*),

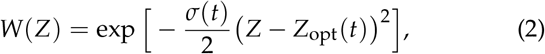

where *σ*(*t*) *>* 0 (the inverse width of the fitness function) measures the selection strength. At time *t* = 0, a sudden environmental change occurs and the trait optimum jumps from the ancestral optimum, 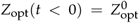, to a new optimum, 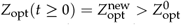. The selection strength, *σ*(*t*), may or may not change at this time and can also vary before or after.

Denote the frequency of the *A*_*i*_ allele in the population as *p*_*i*_. New mutations from *a*_*i*_ to *A*_*i*_ arise at rate *μ*_*i*_ per generation and back-mutations at rate *ν*_*i*_. Loci may be linked, assuming a single linear chromosome with recombination rate *r* between neighboring loci. Prior to the environmental change, population variation segregates at mutation-selection-drift balance. If locus mutation rates are sufficiently small (and/or selection is sufficiently strong), the trait mean is 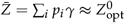. Moreover, allele frequencies, *p*_*i*_, in the SGV follow a U-shaped distribution, such that, for an ancestral trait optimum of 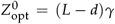, there are *d*≤ *L* loci almost fixed for the *a*_*i*_ allele (*p*_*i* ≈_ 0, “beneficial variation”), while *L* − *d* loci are almost fixed for the *A*_*i*_ allele (*p*_*i* ≈_ 1, “deleterious variation”) at *t* = 0. The case of no SGV is formally included in this framework as the limit of very strong purifying selection (very large *σ*(*t*) for *t <* 0).

After the environmental change, the trait mean 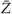 starts to move toward the new optimum. During this adaptation process, we take “snapshots” of the population, *i*.*e*., we record all allele frequencies when the trait mean reaches a threshold of 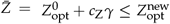 (see Fig. 1). Here, *c*_Z_ measures adaptation at the level of the phenotype in units of mutational steps. The *joint distribution of allele frequencies* across evolutionary replicates at the threshold points serves as our measure of the adaptive architecture (following Höllinger *et al*. 2019; Barghi *et al*. 2020).

**Figure 1.**
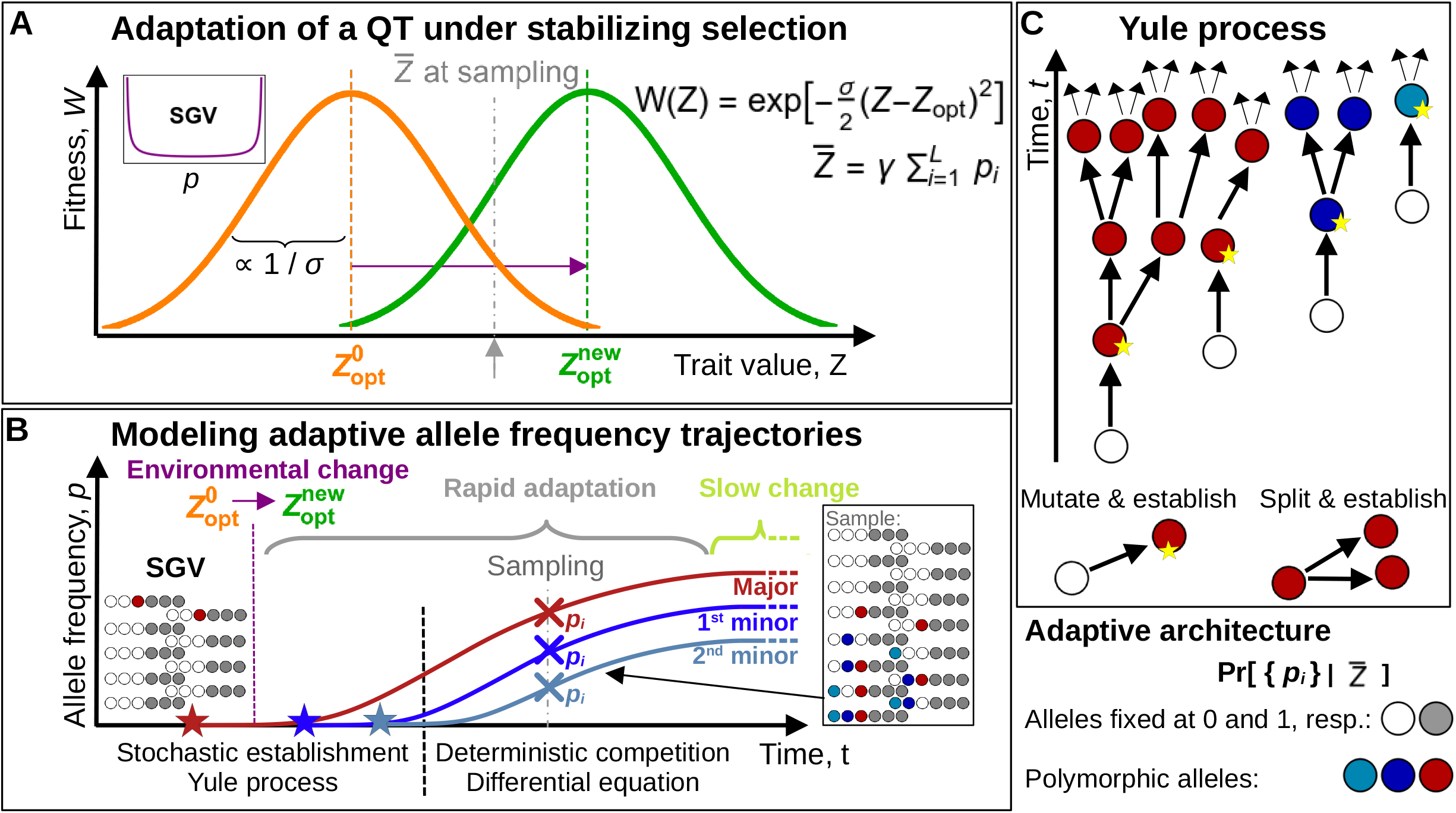
Modeling approach. (A) After an environmental change, the optimum of a trait *Z* under Gaussian stabilizing selection, *W*(*Z*), shifts from 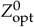 to 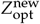 (orange and green bell curves). The population adapts from mutation-selection-drift balance (SGV). Allele frequencies at all *L* loci underlying the trait are recorded once the trait mean, 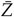, reaches a threshold value (gray arrow). (B) We dissect the adaptive process into separate phases: During the early establishment phase, both prior and after the environmental change, mutant allele frequencies are typically small. Mutants evolve largely independently, but are strongly affected by the stochastic forces of new mutation and genetic drift. Once beneficial mutants have grown to higher frequency, stochasticity can be ignored, but competition and epistatic interaction become relevant. As long as the mean phenotype is not too close to the new optimum, 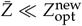, directional selection prevails and leads to rapid adaptation. Close to the optimum, the dynamics slow down, and evolution under weak disruptive selection leads to a depletion of variation. (C) We use a multi-type Yule branching process to track the allele counts at different loci during the establishment phase. The process only follows “immortal” mutant lineages that escape loss due to drift. New immortal lines originate either via new mutation at all loci, or by birth events (splits) of existing lines.

#### Individual-based simulations

We resort to individual-based (IB) simulations to compute the full adaptation dynamics in discrete time for a population of *N*_e_ haploids. Each generation, *N*_e_ mating pairs are chosen via stochastic acceptance (Lipowski and Lipowska 2012) according to their fitness Eq. (2). For each pair, a binomially distributed number of crossover points are randomly placed on a single chromosome to construct the genotype of a single recombinant offspring individual: The probability of a recombination event between neighboring sites is *r*. This is followed by bidirectional mutation (*a*_*i*_ *↔A*_*i*_), where the number of mutated sites in the entire population is Poisson distributed with parameter *N*_e_ *Lμ*. This completes the simulated life cycle.

All individuals are initialized with *d* randomly-chosen loci carrying the allele *a*_*i*_ and *L* − *d* loci with *A*_*i*_ in order to match the ancestral optimum, 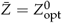. Then, the population equilibrates for 20*N*_e_ generations under constant stabilizing selection to reach mutation-selection-drift balance prior to the environmental change. At *t* = 0, the optimum jumps to 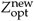, and the population starts to adapt. Allele frequencies at all loci and the full phenotype distribution are recorded when 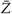 first reaches a prescribed threshold value, 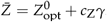. We usually consider a population of size *N*_e_ = 1 000 and summarize results for 10 000 replicates. Further details can be found in the *Computational Appendix* (uses *Mathematica* by Wolfram Research 2019) and the provided IB-simulation script (in *C++*, Stroustrup 2013).

#### Linkage equilibrium simulations

For weak selection and/or strong recombination, we can ignore all linkage disequilibria (LD) and reduce the multi-locus dynamics to the dynamics of single-locus frequencies, *p*_*i*_. Analytic expressions for the single-locus dynamics can be derived for a large class of models (see *Mathematical Appendix*). For Gaussian selection, in particular, the single-locus equations are well-known (Wright 1931; Barton and Bengtsson 1986; de Vladar and Barton 2014; Jain and Stephan 2015, 2017b),

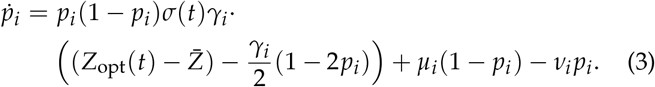

The first selection term, 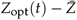, describes directional selection toward *Z*_opt_(*t*). It dominates as long as the trait mean is sufficiently far from the optimum, which is typically the case in the early phase of rapid adaptation. The second term, (*γ*_*i*_ /2)(1 − 2*p*_*i*_) corresponds to disruptive selection and dominates the dynamics once the trait mean gets very close to the new optimum. In this later phase of adaptation, selection for reduced genetic variation drives alleles to the boundaries of the allele frequency range – either to loss or fixation.

We can assess these dynamics using efficient Wright-Fisher simulations that track loci separately in discrete time. Allele frequencies at a locus in the offspring generation are generated by forward and backward mutation with equal rates, Θ_*i*_ /2 = *N*_e_ *μ*, followed by binomial sampling to implement selection and drift. Let the sampling weight of the *a*_*i*_ allele be normalized to 1, then the *A*_*i*_ allele has the weight

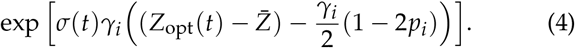

The dynamics at individual loci influence each other via the mean trait, 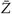, which is recorded in every generation. Allele frequencies at all loci are reported whenever 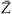 reaches a threshold value.

We usually simulate a population of *N*_e_ = 10 000 haploid individuals, with 3≤ *L* ≤ 10 000, loci underlying the trait, and evaluate 10 000 replicates per parameter combination. Prior to the environmental change, we let the population equilibrate for at least 8*N*_e_ generations to build up SGV. Further details can be found in the *Computational Appendix* and the included LE-simulation script (in *C++*).

### Analytical approximations

To obtain an analytical approximation for the joint allele frequency distribution during rapid adaptation, we extend a framework developed for a binary trait in Höllinger *et al*. (2019) to the case of an additive QT. The approach separates the dynamical process into two phases: an initial stochastic *establishment phase* and a subsequent *deterministic phase* during which established mutants grow and compete (see Fig. 1B). Similar approaches have been used by Uecker and Hermisson (2011) and Martin and Lambert (2015) to describe the dynamics of a single locus and by Götsch and Bürger (2023) for independent loci.

We start with a population in mutation-selection-drift balance and the trait mean, 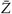, close to its ancestral optimum, 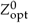. During the first, stochastic phase, mutant frequencies both before and directly after the environmental change are small, such that epistatic interactions can be neglected. Assuming linkage equilibrium (LE), the dynamics at individual loci (due to drift, mutation, and selection) are approximately independent and can be described by a branching process. Once the frequencies of beneficial alleles become sufficiently large, the phenotype adapts rapidly towards the new optimum, 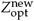. Since drift and new mutation can be ignored during this phase, the allele frequency dynamics are effectively deterministic. They are driven by directional selection, but are no longer independent, because alleles at different loci are coupled due to (fitness) epistasis. Mutants also compete for their relative contribution to adaptive change in the mean trait. The end of this phase is reached, when the rapid change of the mean population phenotype slows down in the vicinity of the new optimum and the dynamics of disruptive selection and drift take over.

#### Yule branching process

During the establishment phase, we describe the allele dynamics by a multi-type *Yule process* (see Fig. 1C). This process tracks the origin and spread of mutations that successfully establish within the population, *i*.*e*., that escape stochastic loss and leave descendants in the population until observation. It thus corresponds to a coalescent genealogy, but is constructed forward in time, at multiple loci simultaneously. Details of the construction are given in the *Mathematical Appendix*.

We denote the establishment probability of a new mutant copy (the probability that descendants still exist at the time of sampling) as *p*_est_(*t*). Such copies found new “immortal lineages” of the Yule tree. Since selection is time-dependent, also *p*_est_(*t*) depends on time. In particular, establishment of a later-beneficial *A*_*i*_ mutant is much less likely if it originates before the environmental change (*t <* 0), while the allele is still deleterious. Analytical approximations for *p*_est_(*t*) lead to complex expressions (Uecker and Hermisson 2011), but this is not needed here. Our results simply exploit the fact that for small *p*_*i*_ the establishment probability of mutants with the same phenotypic effect is (approximately) identical for all loci.

There are two events that create new immortal lines of the Yule process: New successful mutation (before and after the environmental change) seeds novel Yule trees at rate

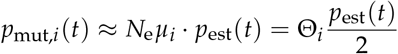

for locus *i*. Birth events lead to splits of existing immortal lines – branching of an existing Yule tree – at rate

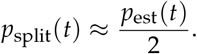

We start the Yule process at some time *t*≤ 0 before the first successful mutation has originated and stop it when a certain number of immortal lineages has been generated. At this point, we assess the distribution of immortal lineages across all loci. Since the rates of all events at loci with equal effect are proportional to the same establishment probability, *p*_est_(*t*), we can drop this common factor if we are only interested in the sequence of events and not in their timing. Mathematically, this corresponds to a time rescaling. On the new timescale, the process is time-homogeneous, with constant rates *p*_mut,*i*_ = Θ_*i*_ and *p*_split_ = 1. For this simple process, we find that the distribution of *ratios* of allele frequencies of beneficial *A*_*i*_ mutants at the end of the stochastic phase is given by an inverted Dirichlet distribution (see Höllinger *et al*. 2019, and the *Mathematical Appendix*).

#### Deterministic phase

Once the allele frequencies are no longer small, and as long as the trait mean is not yet very close to the new optimum, the dynamics of beneficial *A*_*i*_ mutants are well described by the deterministic *directional selection model* (Jain and Stephan 2017b),

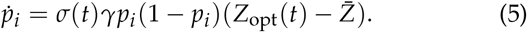

In terms of odds, *u*_*i*_ := *p*_*i*_ /(1 − *p*_*i*_), this reads

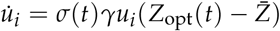

such that odds ratios, *u*_*i*_ /*u*_*j*_, remain constant under the dynamics,

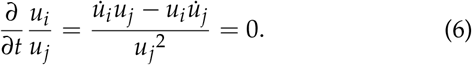

For small allele frequencies, *u*_*i*_ /*u*_*j*_ ≈ *p*_*i*_ /*p*_*j*_. The distribution of odds ratios thus approximately follows the inverted Dirichlet distribution derived above from the Yule process. Due to Eq. (6), the *u*_*i*_ maintain this distribution throughout the deterministic phase. To obtain a result for the joint allele frequency distribution, we need to transform the distribution of the odds *u*_*i*_ back to the frequencies *p*_*i*_ at the stopping condition for the mean trait 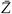.

In a wide parameter region, when effects of rare *a*_*i*_ mutants (that change the trait mean in the opposite direction of phenotypic adaptation) can be ignored, the condition on frequencies *p*_*i*_ and corresponding odds *u*_*i*_ of *A*_*i*_ mutants at the loci 1 to *d* (beneficial variation) at the sampling point reads

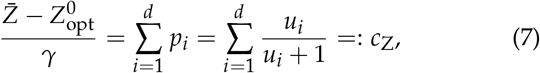

with *c*_Z_ ∈ [0, *d*]. We then obtain the joint distribution of the *p*_*i*_ with 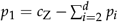 and

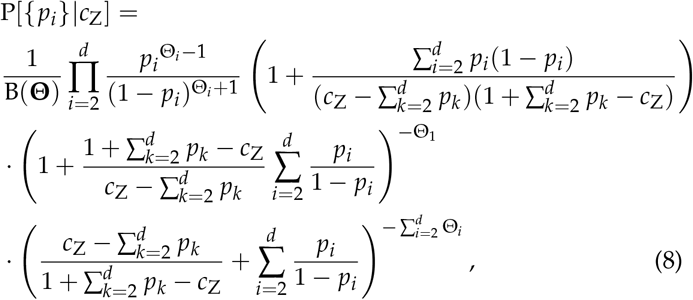

for *i* = 2, …, *d*, where

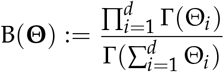

is the multivariate *beta*-function and Γ is the *gamma*-function (see *Mathematical Appendix*). Eq. (8) is the analog of the joint distribution function for adaptation of a bivariate trait (Höllinger *et al*. 2019, Eq. 8). Note that the expression does not depend on any selection parameters (before or after the environmental change), but only on the mutation rates, Θ_*i*_. For two loci, in particular, the marginal distribution at the second locus can be derived explicitly (*p*_2_ = *p*),

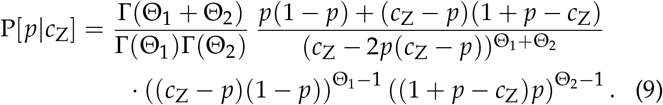

For three or more loci, marginal distributions can only be obtained by (*d* −1)-fold integration. The special functional form of the joint distribution (*cf*. Ghorbel 2009) and its connection to the gamma distribution allow for efficient numerical techniques and precise results even for highly polygenic traits and *d* of the order of 100 or 1 000. This is explained in detail in the *Computational Appendix*.

For high levels of SGV (large Θ_*i*_, many loci) and/or small adaptation distances (small 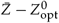 at sampling), the contribution of segregating minority *a*_*i*_ mutants contributing to deleterious variation cannot be neglected. It is still possible to derive (and numerically evaluate) an expression for the joint distribution of allele frequencies at all *L* loci underlying the trait for this case. Since the terms become increasingly complex, the results (and all derivations) are relegated to the *Mathematical Appendix*. The associated, complete numerical solution procedure can be found in the *Computational Appendix*.

## Results

Below, we compare our analytical results with comprehensive computer simulations and discuss characteristic features of the adaptive architecture of a QT across broad parameter ranges in terms of selection strength, mutation rates, number of loci in the genetic basis, linkage, presence/absence of SGV, and distance of the adaptive phenotype from its ancestral value.

Loci have equal strength in our model, and for simplicity we also assume equal locus mutation rates in the results part. This leaves two main sources of differences between loci: asymmetric initial conditions (some loci are nearly fixed for *A*_*i*_ alleles and others for *a*_*i*_ alleles before the environmental change) and the stochastic effects of mutation and genetic drift. We begin by explaining how we define and represent “adaptive architecture”.

### Adaptive architecture

Following Barghi *et al*. (2020), the adaptive architecture of a trait informs about the number and relative contributions of loci that collectively cause a given level of phenotypic adaptation. By definition, the joint allele frequency distribution at all loci in the genetic basis of the trait, taken across replicates at threshold points of phenotypic adaptation, provides an exhaustive description. However, for a trait with a genetic basis of more than a few loci, this is a complex high-dimensional quantity that needs to be projected onto one-dimensional marginal distributions for visualization.

Rather than simply marginalizing for a fixed locus, we follow Höllinger *et al*. (2019) and derive marginal distributions for groups of loci ordered by allele frequency. Therefore, we order all loci according to the frequency of the *A*_*i*_ allele in each evolutionary replicate. Since all locus effects are equal, this also corresponds to their relative contribution to the phenotype. We then construct univariate frequency distributions for loci with the same frequency rank across replicates (which may be different physical loci between replicates). For *L* loci in the genetic basis of the trait, this yields a set of *L* size-ordered marginal distributions to visualize the original *L*-dimensional joint distribution.

In designating frequency-ordered loci, we focus on the loci with *A*_*i*_ as minority allele (the “beneficial variation”), since these contribute the largest allele frequency changes. We therefore often discard the (*L* −*d*) loci with the largest frequency of the *A*_*i*_ allele (assume them to be fixed) and call the locus with the (*L*− *d* + 1)-largest frequency the “major locus” of the adaptive process. Subsequent loci are called the “first”, “second”, “third”, *etc*. minor loci, accordingly. This is consistent with Höllinger *et al*. (2019) and also with standard nomenclature. Indeed, although all locus effects are equal in our model, the major locus would typically yield the strongest signal in an association study of the adaptive trait (if there is any signal at all) because it has the largest frequency change.

#### Sampling in “pheno-time”

The adaptive architecture relates adaptive change at the level of the phenotype to changes in the underlying genotype. Accordingly, we record allele frequencies during the adaptive process not after a fixed time, but when a stopping condition 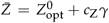 is reached for the mean phenotype (fixed *pheno-time*). For each sampling point, we calculate analytical predictions for the joint distribution of allele frequencies and compare them with numerical simulations.

In this manuscript, we focus entirely on the architecture of the early, rapid phase of adaptation, while the trait is still predominantly subject to directional selection. We thus consider sampling points before the trait mean reaches the new optimum, 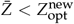. The architecture observed at these points is transient and is eventually replaced by a pattern of adaptive substitutions as stabilizing selection at the new optimum drives alleles to either loss or fixation. In the following sections, we describe the key aspects of this transient architecture for scenarios of increasing complexity.

### Architecture types and background mutation rate

We start with a particularly simple scenario that nevertheless highlights the key features of the basic architecture types. To this end, consider a trait with only three unlinked haploid loci that are all initially fixed for the *a*_*i*_ allele (no SGV). Selection prior to the environmental change is directional toward the lower bound of the phenotype range, 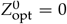. At *t* = 0, the trait optimum switches to the opposite end of the phenotype range, 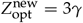. Adaptation occurs from recurrent new mutation, with the same rate Θ = 2*N*_e_ *μ* at all three loci. We record the frequencies of *A*_*i*_ alleles when the trait mean has increased by a single mutational step, 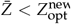 (*i*.*e*., *c*_Z_ = 1), which is two steps below the new optimum.

Fig. 2 shows the marginal distributions of the major locus with the largest allele frequency at sampling (red), followed by the first and second minor loci (dark and light blue, respectively) for various values of the mutation rate, where Θ_bg_ = 2Θ = 4*N*_e_ *μ* measures the *background mutation rate* that is further discussed below. We show Wright-Fisher simulation results (assuming LE) for two models: dots for the full model (Eqs. 3 and 4) and asterisks for the simplified directional selection model, Eq. (5), without the disruptive selection term of the full model (the second term in Eq. 4). The analytical predictions (solid lines) show a perfect fit for the directional selection model, for which they were derived. For the full model (dots) deviations become visible, as all marginal distributions are slightly pushed towards the boundaries of the frequency range by the disruptive selection term (with a relative strength of up to 1/4 of the strength of directional selection). In more detail, we observe the following:

**Figure 2.**
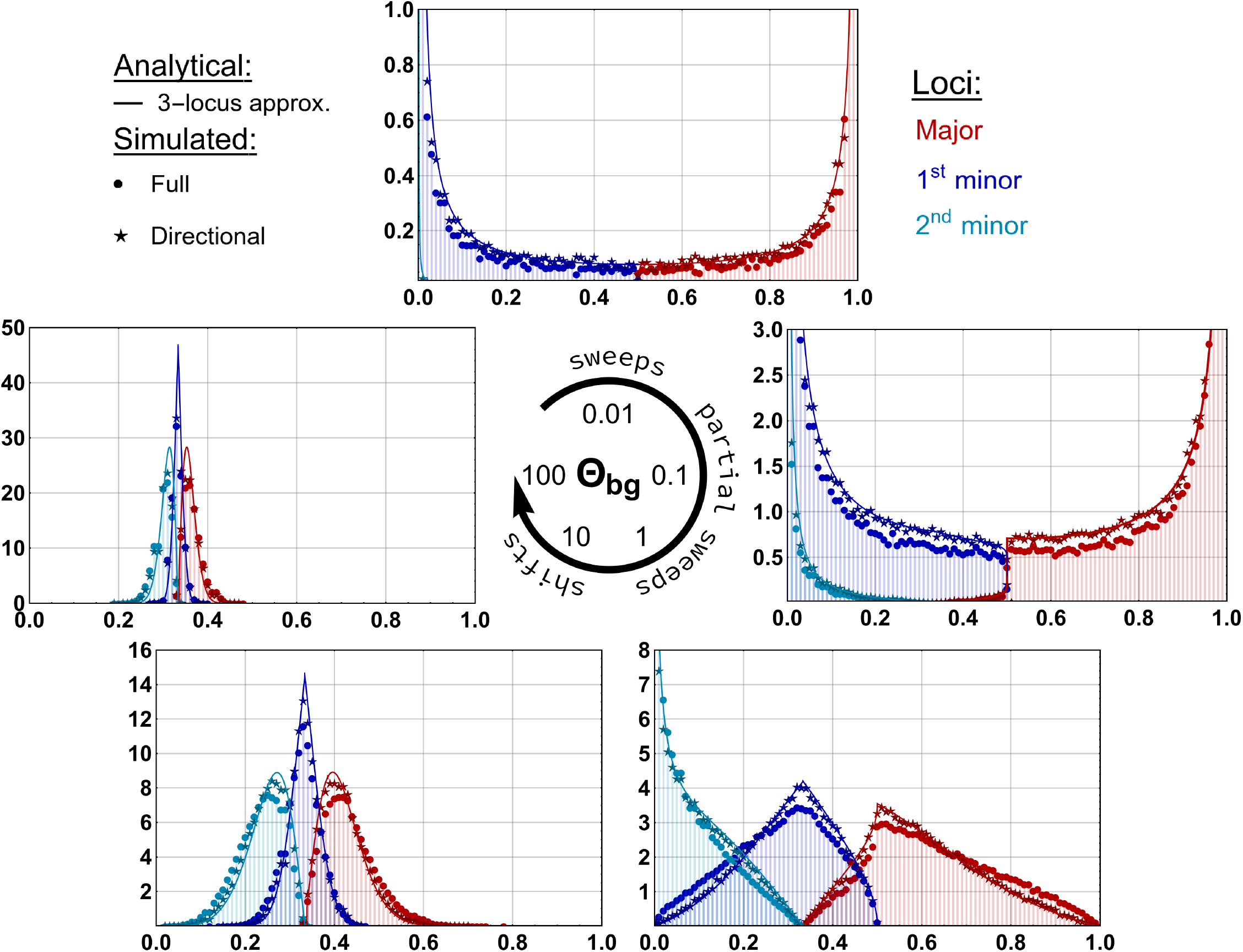
Types of adaptive architecture. After an environmental change, an initially monomorphic trait adapts from *Z* = 0 towards a new optimum at 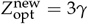, using new beneficial mutations at three loci. We observe the marginal distributions of ordered allelefrequency classes when phenotypic adaptation has proceeded by one mutational step, 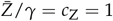. Analytical predictions (lines) are compared to the numerical simulations of the directional selection model (asterisks) and the full model (dots). We find three distinct patterns of adaptive architectures: single selective sweeps for low Θ_bg_, partial sweeps for intermediate Θ_bg_ and a collective, shift-like response of all 3 loci for high Θ_bg_. *N*_e_ = 10 000, *N*_e_ *σγ*^2^ = 100, 100 000 replicates. Note the differences in the y-axes ranges.

- For very low mutation rates, Θ_bg_ ≲ 0.1, the adaptive change in phenotype is usually accomplished by a single locus. Adaptation thus occurs by a classic selective sweep at the first locus where an *A*_*i*_ allele appears and is picked up by selection. This locus is the major locus with a pronounced peak of its frequency distribution (red) at 1. Both minor loci hardly contribute to adaptation at all (blue distributions peak at 0). Note that this strong heterogeneity across loci is not visible in the marginal distribution of a single focal locus, since all three loci can end up as the major locus with equal probability.
- For intermediate mutation rates, 0.1 ≲ Θ_bg_ ≲ 10, the distributions of major and minor locus frequencies are still clearly distinct. However, minor loci now also contribute significantly to phenotypic adaptation, leading to two (Θ_bg_ = 0.1: major and first minor) or even three (Θ_bg_ ≥1: all loci involved), partial sweeps.
- Finally, at high mutation rates, Θ_bg_ 2 10, the three marginal allele frequency distributions of major and minor loci gradually converge. Phenotypic adaptation occurs via the collective and homogeneous response at all three underlying loci (frequency shifts around 1/3 at all three loci for a 3-locus trait).

The striking differences in adaptive architecture, from heterogeneous single sweeps to homogeneous collective shifts, reflect the decreasing role of stochasticity for the evolutionary process as Θ_bg_ increases. All characteristic features are captured by the analytical approximation of the joint distribution, Eq. (8).

#### The background mutation rate Θ_bg_

The type of architecture, between sweeps and shifts, is already decided early in the initial stochastic phase of the adaptive process. As it turns out, its degree of homogeneity is largely determined by a single composite parameter, the so-called background mutation rate Θ_bg_ (Höllinger *et al*. 2019). For a haploid model with equal locus mutation rates *μ* and *d* ≤*L* loci carrying an *a*_*i*_ majority allele for *t <* 0, it reads

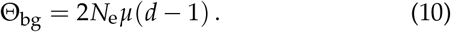

Θ_bg_ can be understood as a measure of redundancy (Barghi *et al*. 2020), because it measures the total mutational input of beneficial alleles with an equivalent effect to the allele at the major locus and are thus redundant options for the next adaptive step. Its role in the process can also be understood as follows. Consider the early stochastic phase of adaptation described by the Yule process. At some point in time, the first Yule tree is seeded by the first *A*_*i*_ mutation that escapes stochastic loss. Then 1/Θ_bg_ is the average waiting time for a second Yule tree to emerge at a different locus (measured on the timescale of the exponential growth of the first Yule tree, which has a split rate of 1). Θ_bg_ thus quantifies the expected head start of the frontrunner allele over its competitors, and thus the heterogeneity of allelic contributions to the adaptive trait.

### Effect of selection, trait basis size, and SGV

Equipped with these concepts, we can explore the adaptive architecture of an additive QT across a wider parameter space. In particular, we study the effects of the size, *L*, of the genetic basis and the selection strength, before and after the environmental change. We also include SGV as a source of adaptive variants.

Our basic setup is as follows: for a trait of size *L* (we use *L* = 10 and *L* = 100 in the figures), we place the ancestral optimum in the middle of the phenotype range, 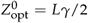, where we let the population equilibrate to mutation-selection-drift balance. LE is assumed. At time *t* = 0, the trait optimum jumps to a new value at a distance of three mutational steps, 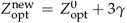. As in the previous section, we record the adaptive architecture once the trait mean has increased by one mutational step (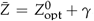, *i*.*e*., *c*_Z_ = 1). Adaptation over larger distances and effects of LD are discussed in the following sections. When comparing adaptive architectures of traits with different size *L* of their genetic bases, we need to decide how model parameters are scaled. Following the insights of the previous section, we first make sure that the background mutation rate Θ_bg_ = 2*N*_e_ *μ*(*L*/2 − 1) is kept constant in all comparisons. We further keep the scaled selection strength *N*_e_ *σγ*^2^ prior to the environmental change constant, which leads to total expected levels of SGV that are (almost) independent of *N*_e_ and *L* (see the *Mathematical Appendix*, Remark 3, for details).

Fig. 3 summarizes the effects of three parameters on the adaptive architecture: the background mutation rate Θ_bg_ (between 0.01 and 100), the selection strength *N*_e_ *σγ*^2^ (10 for weak and 100 for strong) and the number of loci *L* (10 and 100). The simulation results are compared with analytical predictions. Most importantly, we see that neither selection strength nor trait size nor the origin of the mutations from SGV or new mutation has a qualitative effect on the adaptive architecture (compare panels in the same row of Fig. 3 and with the same Θ_bg_ in Fig. 2). As in the three-locus case, the background mutation rate emerges as the (only) crucial parameter for determining the architecture type or mode of adaptation: For Θ_bg ≲_ 0.1 we observe a single sweep at the major locus, while all other loci remain fixed at *p*_*i*_ = 0 or 1. At intermediate values, 0.1 ≲ Θ_bg_ ≲ 10, adaptation proceeds via partial sweeps at a limited number of loci. The relative contribution of different (but identical) loci remains very heterogeneous (major and minor loci), highlighting a dominant role of genetic drift. For large Θ_bg_ ≥ 0, very many loci contribute to adaptation. Individual contributions become more homogeneous, and thus approach 1/*L*, leading to “subtle allele frequency shifts” for traits with a large genetic basis. For a detailed, quantitative assessment, we distinguish parameter regions where the purging of deleterious mutations does or does not play a role for phenotypic adaptation.

**Figure 3.**
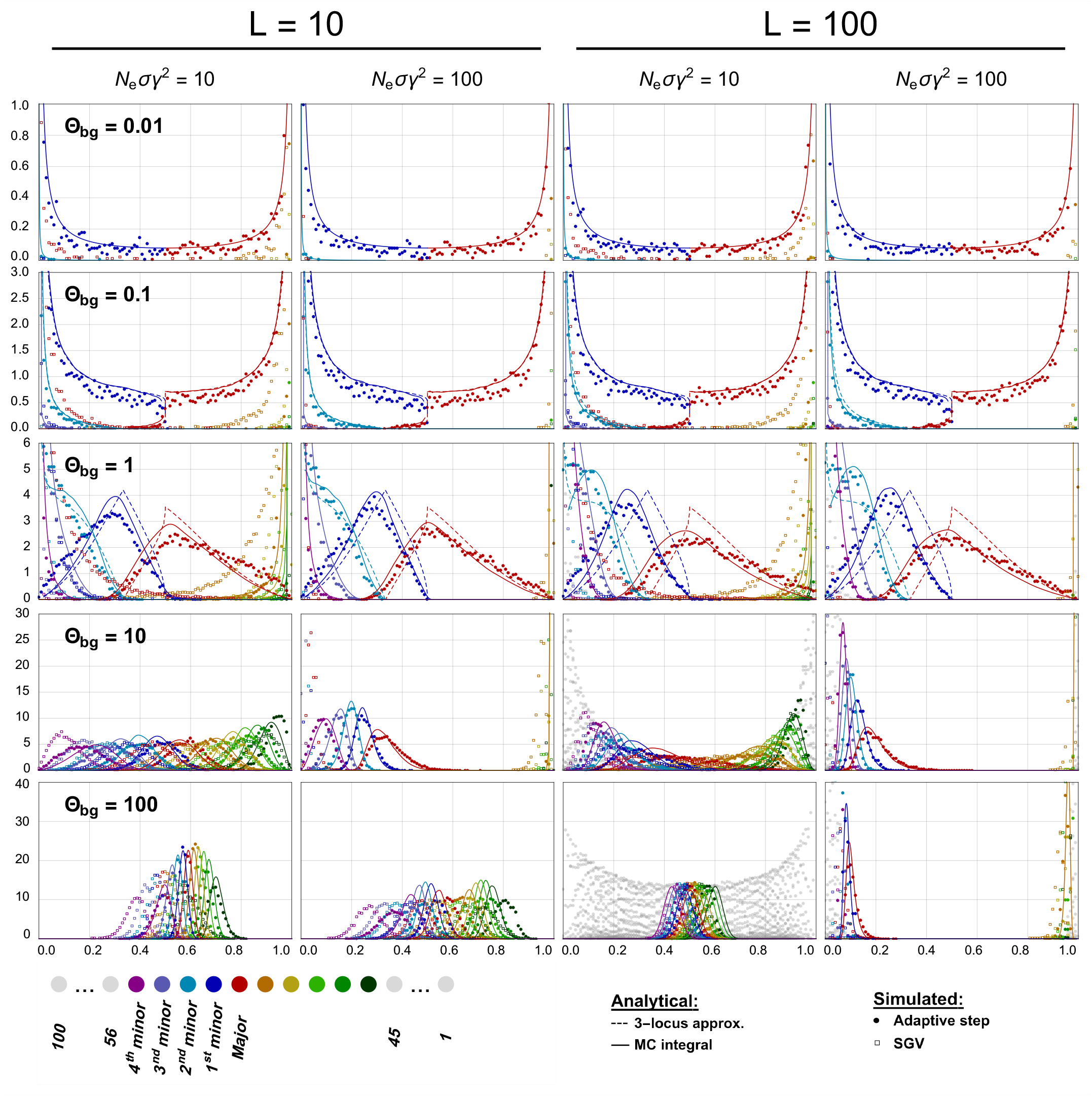
Effect of selection strength and trait basis size on the adaptive architecture of a QT. The shape of the adaptive architecture, indicating selective sweeps (Θ_bg_≤ 0.01), partial sweeps (0.1 ≤Θ_bg_ *<* 10), or shifts (Θ_bg_ ≥10), is independent of the selection strength and the number of loci in the trait basis. For a trait of size *L* = 10 or *L* = 100, the optimum shifts at time *t* = 0 from 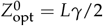 to 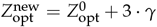. Stabilizing selection is either weak (*N*_e_ *σγ*^2^ = 10) or strong (*N*_e_ *σγ*^2^ = 100). The allele frequency distributions of all loci ordered according to their frequency are obtained from simulations at *t* = 0 (SGV, open symbols) and after adaptation of a single mutational step, 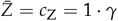 (closed symbols). Lines show the analytical approximation, using either the full *L*-locus model (solid lines) or a 2- or 3-locus formalism where closed expressions can be derived (dashed lines), see the text for details. For *L* = 100, the distributions of the 10 “middle” loci (46 through 55), which correspond most closely to the 10-locus case, are shown in color. All other distributions are shown in grey. *N*_e_ = 10 000, 10 000 replicates. Note the differences in scaling of the y-axis.

#### Adaptation dominated by beneficial variants

When a trait under stabilizing selection adapts to a shift in the optimum, there are two ways how selection on SGV can contribute: either by increasing the frequency of beneficial mutants (*A*_*i*_), or by eliminating deleterious mutants (*a*_*i*_) that change the phenotype in the opposite direction. In our example, with the original trait optimum in the middle of the phenotype range, half of the loci may carry rare *A*_*i*_ alleles and the other half rare *a*_*i*_ mutants for *t*≤ 0. After the environmental change, the former constitute the “beneficial variation” and the latter the “deleterious variation”. There is a large parameter range in which the maximal contribution of all rare deleterious variants to the change in 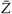 (by eliminating all of them) is small relative to the total adaptive change in 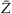. This is the case if mutation rates are small and/or stabilizing selection is strong, such that levels of SGV are low. In contrast, even single beneficial mutants can make a substantial contribution if they progress to fixation from a low starting frequency or if they originate as a new mutation after the environmental change.

In Fig. 3, there is hardly any deleterious variation segregating at sampling for weak selection and Θ_bg_≤ 0.1, and for strong selection and Θ_bg_ ≤1 (orange/yellow/green colors). We can thus assume that the corresponding loci are fixed for the *A*_*i*_ allele in the joint distribution and use Eq. (8) to describe the joint variation at the beneficial loci. The analytical prediction is independent of selection parameters. Its good fit shows that the adaptive architecture, in this parameter region, neither depends on the strength of stabilizing selection prior to the environmental change (and, hence, on the amount of SGV) nor on the strength of directional selection that drives the adaptation.

#### Contribution of deleterious variation

For high initial levels of SGV (high mutation rates and/or weak selection), deleterious variants are not yet fully purged at the sampling point (orange/yellow/green distributions in Fig. 3). Since the beneficial variation must compensate for the effect of the deleterious variation, this affects the distributions at all loci. In particular, this makes the adaptive architecture dependent on selection *before the environmental change*, because the strength of stabilizing selection affects the amount of SGV. As an example, consider the case *L* = 10 and Θ_bg_ = 1 in Fig. 3. With weak selection, substantial amounts of SGV accumulate before the environmental change (shown as open squares), and some of the deleterious variation is not yet eliminated at the time of sampling. In contrast, with strong selection, the deleterious variation is completely lost. The distributions at the beneficial loci are similar in both cases, but not identical. The differences between weak and strong selection become larger for higher mutation rates, such as Θ_bg_≥ 10.

Our approximation can account for the contribution of both the beneficial and the deleterious variation to the adaptive architecture, see the *Mathematical* and *Computational* appendices for details. In short, the joint distribution of beneficial and deleterious minority alleles still follows a transformed Dirichlet distribution, as in Eq. (8), but with an additional parameter *κ* to scale down the frequencies of the deleterious variants relative to the beneficial variants. This scaling factor depends on the total amount of SGV and thus also on the selection intensity before the environmental change. In the figure, we use this extended approximation in all panels with Θ_bg_ ≥1, where non-trivial contributions of deleterious variation can occur. For Θ_bg_≤ 0.1, the scaling factor becomes *κ*≈ 0 and the extended approximation reduces to Eq. (8).

#### Effect of the number of loci L underlying the trait

Even if a trait is highly polygenic, phenotypic adaptation can be oligogenic if the number of loci that contribute to the adaptive change is much smaller. Indeed, our results show that the polymorphic part of the adaptive architecture is hardly affected by the size of the genetic basis at all, as long as *L* is sufficiently large (or the background mutation rate Θ_bg_ sufficiently small) such that typically fewer than *L* loci are polymorphic. In this case, differences in the number of loci *L* only lead to differences in the number of fixed loci at frequency 0 or 1 (compare *L* = 10 and *L* = 100 in Fig. 3 for Θ_bg_≤ 0.1 and approximately still for Θ_bg_ = 1). This necessarily changes for larger background mutation rates once x A theory of oligogenic adaptation of a quantitative trait traits with a smaller genetic basis run out of further loci that could contribute to adaptation (Θ_bg_ ≥10 in Fig. 3).

We can use the approximate invariance of the adaptive architecture on *L* to describe the adaptive dynamics of a trait with a large genetic basis by a simpler model with few loci and rescaled locus mutation rates. As explained above, 1/Θ_bg_ is the expected waiting time between the origin of the first and second beneficial mutation that contribute to phenotypic adaptation (the first and second Yule tree in our framework). Following Höllinger *et al*. (2019), we can refine this approach and match the waiting time between the first Yule tree and its *j*th follower to approximate the distribution of the *j*th minor locus. Details are given in the *Computational Appendix*.

In Fig. 3, we show how the marginal distributions of the loci with the largest contribution to the adaptive change for traits with *L* = 10 or 100 can be approximated by a 2-locus model (for Θ_bg_ = 0.01) or a 3-locus model (for Θ_bg_ = 0.1 and 1). The match is excellent (with lines almost indistinguishable from the full approximation for Θ_bg_ ≤0.1) whenever the number of contributing loci does not exceed the size of the approximating model. For Θ_bg_ = 1, where often more than 3 loci contribute, deviations appear. In particular, oligogenic adaptation beyond single sweeps is a collective phenomenon that cannot be reduced to a single-locus picture. Due to our conditioning on the phenotype (and also due to fitness epistasis), the shapes of the marginal distributions at different loci with segregating alleles are not independent.

#### Limits of the analytical approximation

The approximation produces a good fit of the simulation data, as long as the relevant alleles are confined to relatively small frequencies ≲ 10% in the SGV. When the initial allele frequency distributions (open squares) extend to intermediate frequencies, two factors become important that are not included in the Yule formalism. First, disruptive selection (the term ∼2*p* −1 in Eq. 3) adds a frequencydependent component that increases the distance between the major and minor locus distributions in the SGV and, consequently, also in the adaptive architecture. This is clearly visible, *e*.*g*., for Θ_bg_ = 10 and weak selection in Fig. 3. Second, for large mutation rates, back-mutation becomes an important factor in shaping allele frequency distributions in the SGV. When strong mutation overwhelms selection, all allele frequencies are pushed toward the mutation equilibrium (*p*_*i*_ = 0.5 for equal forward and backward rates). This decreases rather than increases the distance between major and minor locus distributions. Sometimes the effects of mutation and disruptive selection can almost cancel (*e*.*g*., for *L* = 10, Θ_bg_ = 100, weak selection in Fig. 3), but usually they do not, leading to deviations from the theoretical predictions. Note that high mutation rates *per se* do not compromise the approximation if the SGV is controlled by strong selection (*L* = 100, Θ_bg_ = 100) or if adaptation occurs only from new mutation (*cf*. Fig. 2).

### Adaptation dynamics across larger distances

So far, we have analyzed the types of adaptive architecture that emerge at a single threshold in the very early phase of phenotypic adaptation – a change of 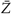 by one mutational step. We now complement this “single snapshot” with a dynamic approach to explore the changes in the adaptive architecture as the mean trait evolves across larger phenotypic distances toward a more distant optimum.

In Fig. 4, we track the adaptation dynamics of a haploid trait with 10 identical loci evolving from mutation-selection-drift balance around an ancestral optimum at 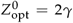 to a new optimum six mutation steps away, 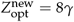. We display marginals of the joint allele frequency distribution at five points throughout the rapid adaptive phase, when the trait mean reaches consecutive threshold values one to five mutational steps away from the ancestral optimum (*i*.*e*., 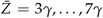). As in the previous section, we examine five orders of magnitude for the background mutation rate, Θ_bg_ = 7 *·* 2*N*_e_ *μ* = 0.01, …, 100. We assume strong selection (*N*_e_ *σγ*^2^ = 100). Additional figures for traits with 10 and 100 loci are provided in the Appendix (Section Adaptation with LE).

**Figure 4.**
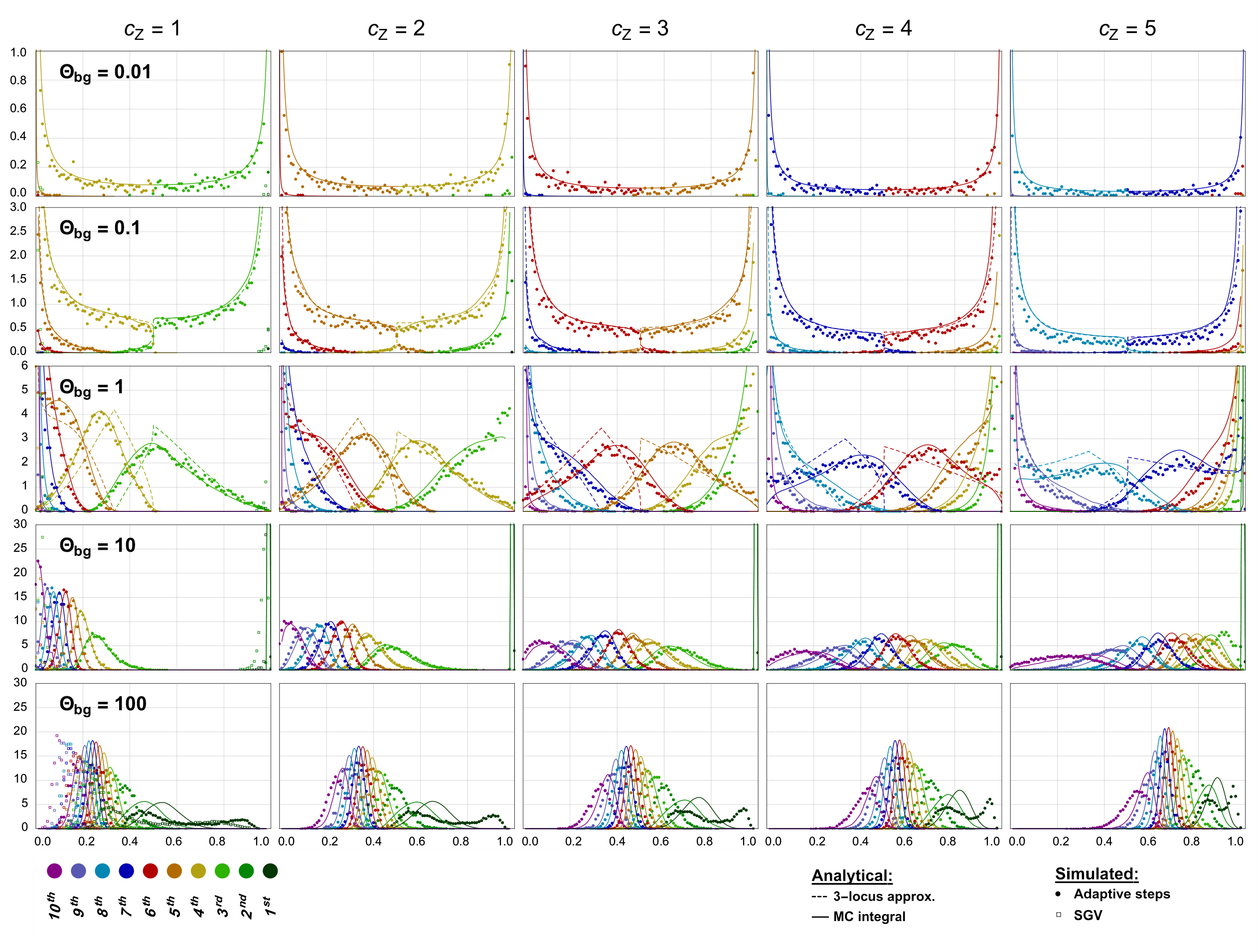
Adaptation across larger distances. Successive snapshots of the adaptive architecture are shown over the course of adaptation of a QT with 10 loci. Populations evolve from mutation-selection-drift balance around the initial optimum 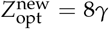 toward a new optimum 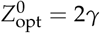 after a change in the environment. Closed symbols show simulation results for the ordered marginal allele frequency distributions after a change in the trait mean 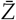 by *c*_Z_ = 1 to *c*_Z_ = 5 mutational steps. The first column also shows the SGV distributions (open squares). Solid lines are analytical predictions for the full model, dashed lines (for Θ_bg_ ≤ 0.1) predictions from an adjusted 3-locus model (see main text). *N*_e_ = 10 000, *N*_e_ *σγ*^2^ = 100, 10 000 replicates. Note the differences in scaling of the y-axis.

The first column in Fig. 4 (after a single mutational step) is analogous to Fig. 3 and shows different types of adaptive architectures, from a single sweep to collective shifts, as the background mutation rate increases. Further columns show that the type of the adaptive architecture is largely preserved at later threshold points, while the color coded distributions of frequency-ordered loci move towards fixation at frequency *p* = 1. For small Θ_bg_ *<* 1, we obtain a pattern typical of mutation-limited adaptation with successive and largely non-interfering selective sweeps. For large Θ_bg_ ≥1, we observe concerted movement of alleles at several (or all) loci simultaneously.

#### Limit shapes of adaptive architecture

From the figure panels of Fig. 4 (clearest for small mutation rates), it appears that the adaptive architecture, when observed at intervals of *kγ* (full mutational steps), approaches a “quasi-stable” limit as phenotypic adaptation progresses. The shape of the joint distribution remains almost invariant, while the role of each locus within this joint distribution changes: a locus that contributes a small-frequency allele after the first step will contribute a larger frequency after the second and later steps, *etc*. In combination, these changes effectively lead to one fewer locus fixed at 0 and one more fixed at 1 with each step, but a constant pattern in the interior of the frequency space.

A stable limit shape requires that sufficiently many loci have traversed the entire frequency range from 0 to 1. This happens quickly in the sweep regime with only few segregating loci, but takes much longer for large Θ_bg_ when adaptation is achieved by small shifts at many loci. A limit is never reached if the distance to the new optimum is too small, or if the trait runs out of further loci that could start their adaptive course at *p* = 0 before the first loci have reached frequency *p* = 1. In our example of a trait with only 10 loci, this is already the case for Θ_bg_ = 10. For traits with a larger genetic basis that adapt to a distant optimum, convergence can also be observed for larger background mutation rates (see the case Θ_bg_ = 10 for a trait with *L* = 100 loci in the Appendix, Fig. S1, and also Fig. 5 for *L* = 10 000 below).

**Figure 5.**
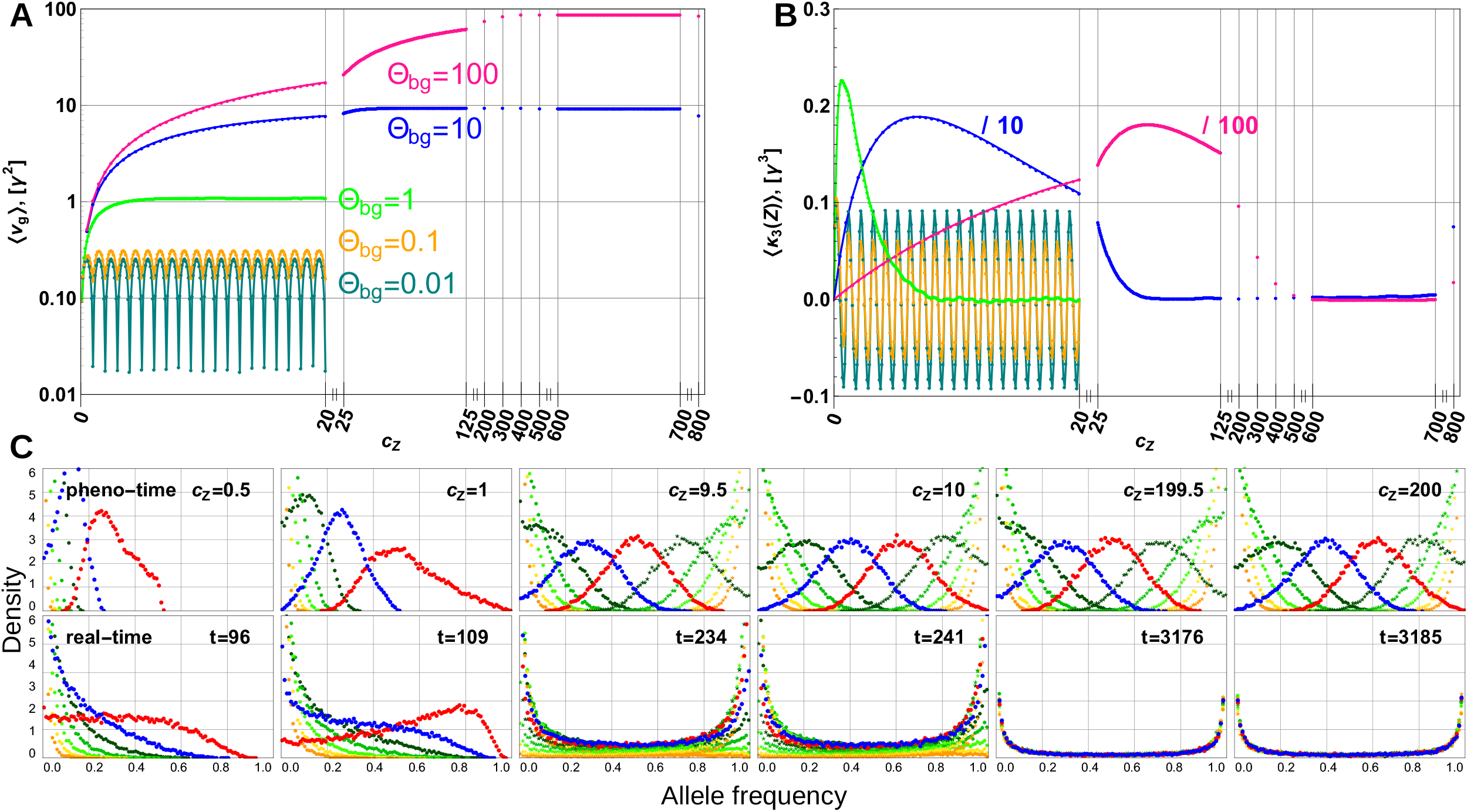
Dynamics of adaptive architecture of a highly-polygenic QT. Panels (A) and (B) show the expected genetic variance, ⟨*v*g ⟩, and skew, ⟨*κ*_3_(*Z*) ⟩, of the trait at sampling points in “pheno-time” for five values of Θ_bg_ = 0.01. Panel (C) shows the adaptive architecture (ordered marginal distributions) for Θ_bg_ = 1, for sampling points in pheno-time (top row) and in real time (bottom). Stopping times are chosen such that the expected mean phenotype 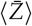 across replicates in real time corresponds to the trait mean 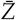(in each replicate run) in the corresponding panel in pheno-time. Two central marginal distributions are shown in red and blue. To highlight the stability of the adaptive architecture at both full and half mutational steps in pheno-time, the colors are not attached to a fixed frequency rank (as in Fig. 4), but to the role of the locus in the joint distribution, *i*.*e*., “red” is the largest locus in the first two panels, the 10th largest for panel 3 and 4, and the 200th largest for panels 5 and 6. Adaptation proceeds from a monomorphic starting state at 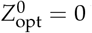 by recurrent, new mutations towards 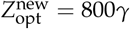. *N*_e_ = 10 000, *L* = 10 000, *N*_e_ *σγ*^2^ = 1, 35 000 replicates. Note that the skew for Θ_bg_ = 10 and 100 in panel (B) is downscaled as indicated: the inset comment “/10” indicates that presented data is 1/10th of the actual data.

A more detailed look shows that the limit distribution in our model is only *quasi*-stable. Indeed, due to the dwindling supply of adaptive material, the expected waiting time between the origin of one beneficial mutation and the next, and thus the distance between the corresponding marginal distributions, increases with each step. As a consequence, the joint distribution gradually becomes more U-shaped (or sweep-like), with lower probability weights at intermediate frequencies, as phenotypic adaptation progresses. In Fig. 4, this is most clearly visible for Θ_bg_ = 0.1 (see also the cases Θ_bg_ = 1 and Θ_bg_ = 10 in Fig. S1). Furthermore, see Fig. S2 in the Appendix for the dynamics of trait summary statistics for *L* = 10 and 100.

We can describe this effect as a reduction in the “effective background mutation rate” that occurs when a trait with finitely many loci adapts over a longer distance. For the final adaptive step to the sampling point at 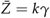 in Fig. 4, there are (10 *k*) “redundant” loci that are not required to reach the threshold, but can still contribute to adaptation and compete with the first *k* loci. We thus have an effective background rate of Θ_bg,eff_ = Θ_bg_(10 −*k*)/7 in terms of the initial background mutation rate (for the first step, where *k* = 3). In our approximation of the 10-locus architecture by an effective 3-locus model (for Θ_bg_ ≤1 in Fig. 4), we match this effective rate Θ_bg,eff_ to obtain a good fit (for Θ_bg_ ≤ 0.1 the differences are hardly visible).

#### Highly polygenic traits

A model of a QT with a constant supply of new mutations (infinite loci or infinite alleles) would lead to a proper limit in the joint distribution without reduction in Θ_bg,eff_. Such “clean” results can also be observed for a trait with a finite, but very large genetic basis. Fig. 5 shows results for adaptation over a large phenotypic distance (up to 800 mutational steps) of a trait with an even much larger genetic basis of *L* = 10 000 loci.

Panels A and B of Fig. 5 show the (scaled) trait variance *v*_g_ = ∑_*i*_ *p*_*i*_ (1 −*p*_*i*_), and the skew *κ*_3_(*Z*) = ∑_*i*_ *p*_*i*_ (1− *p*_*i*_)(1 −2*p*_*i*_), which are both summary statistics of the joint allele frequency distribution (the second and third cumulant). We evaluate these measures at multiple threshold points for short-range (up to *c*_Z_ = 20 mutational steps), mid-range (up to *c*_Z_ = 125) and long-range (up to *c*_Z_ = 800) phenotypic adaptation. For shortrange, in particular, this includes sampling points at non-integer intervals of mutational steps, showing that the joint distribution oscillates with period *γ*. Periodic fluctuations are pronounced for low Θ_bg_, where single peaks in the marginal distributions (representing ongoing sweeps) traverse the frequency space, but become weak and almost disappear in the collective frequency-shifts regime.

While visual convergence to a stable limit period occurs within a few steps for small Θ_bg_ *<* 1, the adaptation distance that is required rapidly increases (roughly proportional to the number of co-segregating alleles) for more polygenic architectures: 9-10 steps for Θ_bg_ = 1, ≈ 70 for Θ_bg_ = 10, and almost 600 for Θ_bg_ = 100. Due to the large size, *L* = 10 000, of the genetic basis, the supply of further beneficial alleles does not change much, even over these distances. As a consequence, the slow change in the adaptive architecture that we have seen for *L* = 10 is no longer visible in the figure. In the Appendix (Section Dynamics of summary statistics), we show how the limit values for *v*_g_ relate to analytical approximations. Convergence of the skewness to (a period around) zero reflects the fact that the limit architecture (if averaged over a period) is approximately symmetric for all values of the background mutation rate.

As shown in panel C of Fig. 5, a stable shape of the adaptive architecture results only for sampling points at constant phenotype (*i*.*e*., in “pheno-time”), but not in real time. Indeed, stochastic effects on the waiting times between successive new mutations lead to a broadening of the marginal distributions, when sampling occurs at a fixed time after the environmental change. For larger times, the probability density piles up at 0 and 1, and all characteristic features of the joint distribution are eliminated.

Our analytical approximation describes not only the limit distribution, but all dynamical changes in the adaptive architecture, as long as selection is primarily directional (Figs. 4 and S1). In particular, this includes a switch from using alleles from the SGV in the initial steps to primarily alleles that enter the population as new mutations in the later adaptive steps. This transition is seamless and occurs without any discontinuity in the shape of the joint distribution. Likewise, the shape of the limit architecture is not affected by the decrease in the strength of directional selection as the trait mean approaches the new optimum.

Once the trait mean closes in on the new phenotypic optimum, the shape of the joint allele frequency distribution eventually does change due to the action of disruptive selection against standing variation. In Figs. 4 and 5 these changes become visible at the last observation point. When we follow adaptation further, selection against genetic variation pushes all frequencies toward the boundaries until all frequency distributions once again assume the typical U-shape of mutation-stabilizing-selection balance. Thus, the shapes of adaptive architecture described above are only transient, their characteristic differences eventually disappearing as the population equilibrates at the new optimum.

### Linkage and ploidy

Our analytical results and all simulations so far assume LE between all loci under selection. This assumption becomes unrealistic for highly polygenic traits when beneficial alleles co-segregate at many loci. In Fig. 6 and in the Appendix (Section Adaptation with linkage), we present IB simulation results to address this issue. In addition, we analyze how our results apply to diploid genetics.

**Figure 6.**
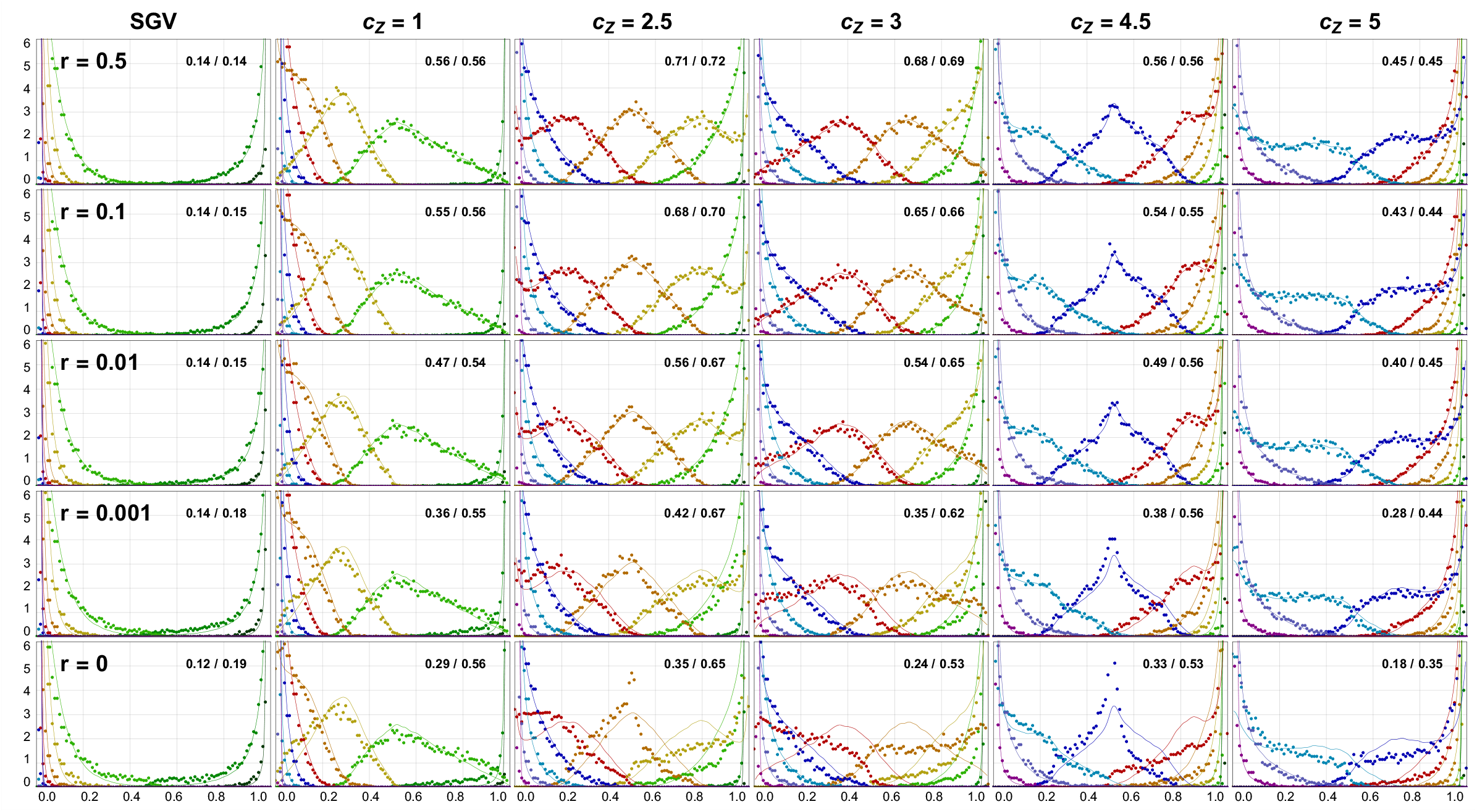
Effect of linkage on the adaptive architecture of a haploid QT (Θ_bg_ = 1). IB simulations with linkage (dots) are compared with LE results (lines). Numbers in the top right corner of each panel show the average genetic variance, *v*_g_ = Var[∑_*i*_ *p*_*i*_], and genic variance, 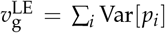 (in units of [*γ*^2^]). While negative LD build up once recombination is weaker than selection (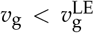 for *r ≲* 0.01 = *σγ*^2^), larger deviations of the adaptive architecture only emerge for (almost) complete linkage. *N*_e_ = 1 000, *L* = 10, *N*_e_ *σγ*^2^ = 10, 10 000 replicates (IB), 125 000 replicates and very mild spline-smoothing for LE simulations.

We use rescaled mutation and selection parameters for our IB simulations with *N*_e_ = 1 000 to match the LE simulations with *N*_e_ = 10 000 of the previous sections. In particular, the selection intensity of *σγ*^2^ = 0.01 prior to the environmental change (that we will use throughout) corresponds to the case of “weak” selection (*N*_e_ *σγ*^2^ = 10) in the previous figures. We assume that the loci underlying the trait are equally spaced on a single, linear chromosome, with recombination probability *r* ≤ 0.5 between adjacent loci. Accordingly, the recombination probability between two loci at distance Δ*ℓ · r* Morgan reads 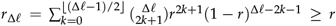. For 10 loci and *r* = 0.01, in particular, 0.01 ≤*r*_Δ*ℓ*_≤ *r*_9_ ≈0.08. This is of the same order as the selection strength, which is *σγ*^2^ = 0.01 prior to the environmental change and 6*· σγ*^2^ = 0.06 directly after the change. Recombination operates faster than selection (*loose linkage*) for *r*≥ 0.1, whereas linkage is *tight* for *r*≤ 0.001.

Fig. 6 shows the effects of linkage on the adaptive architecture of a 10-locus trait with background mutation rate Θ_bg_ = 1. At the environmental change, the trait optimum switches from 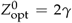 to 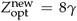. Adaptation starts from SGV and is assessed at several threshold values for 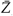, representing both full and half mutational steps. The top row represents the case of free linkage (*r* = 0.5). To validate our simulations and connect them to the previous sections, we also performed LE simulations for the same scenario, which are included in the figure as thin solid lines. For *r* = 0.5, we observe a perfect match for all sampling points, including at 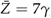, close to the new optimum, where disruptive selection already has a strong effect. The same holds for *r* = 0.1 (loose linkage, second row). Even for *r* = 0.01 and *r* = 0.001 only minor deviations are visible, although linkage is strong and a comparison of the genetic variance, *v*_g_ = Var[∑_*i*_ *p*_*i*_], and genic (or LE-)variance, 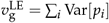 (upper right corner of the figure panels) shows that negative LD build up. Negative LD is expected, both due to negative fitness epistasis and Hill-Robertson interference (Hill and Robertson 1966).

Larger deviations appear only for complete linkage (*r* = 0, bottom row), as a consequence of strong clonal interference and “mutational stacking” (Gerrish and Lenski 1998; Desai and Fisher 2007). Although the dynamics in this limit are entirely driven by the competition among haplotypes, and site frequencies are only a by-product, the qualitative features of the adaptive architecture (as measured by the frequency-ordered single-locus marginal distributions) remain surprisingly robust.

Results for other values of the background mutation rate Θ_bg_ are presented in the Appendix (Section Adaptation with linkage). Generally, linkage effects are very weak for small Θ_bg_ ≪ 1, because multi-locus polymorphism is rare. They also become weaker for (very) large Θ_bg_ ≳ *L*, because recurrent new mutation at a locus reduces LD. Last, not least, we extend our results to diploid genetics. In the absence of dominance, we find perfect agreement (see Fig. S7) with the haploid predictions when the key model parameters (mutation rate, selection strength, effect sizes) are scaled appropriately, see the Appendix (Section Diploids with linkage) for details.

## Discussion

How genetically complex traits adapt to novel environments is a classical question of quantitative genetics. In the current study, we consider the standard model of quantitative genetics, an additive trait under Gaussian stabilizing selection that adapts to a shift in the trait optimum. However, we use a modeling approach (following Höllinger *et al*. 2019) rooted in population genetics. While we measure adaptive progress at the level of the phenotype, we set up an analytical framework to explicitly track the joint allele frequency dynamics at all loci of the genetic basis. The fundamental question is: How does a given level of phenotypic adaptation (change in the trait mean) translate into contributions (allele frequency changes) at the underlying loci? We call this collective genotypic pattern, conditional on the phenotype, the corresponding “adaptive architecture”. Mathematically, it is described by the joint distribution of beneficial allele frequencies across all loci of the genetic basis. Depending on the model parameters, this distribution may describe large allele frequency changes at a few loci (*i*.*e*., sweeps) or small shifts at very many loci. Importantly, it sheds light on a large parameter range in between, where many, but not extremely many loci contribute to what we call *oligogenic adaptation* of a QT.

### Adaptive architectures of a QT

When a QT adapts to a new optimum, there are two phases of adaptive evolution. First comes a (relatively) rapid phase driven by directional selection, during which the trait mean approaches the new optimum. This is followed by a (relatively much longer) equilibration phase of fine-tuning and allele sorting, in which selection against genetic variation and other, weaker evolutionary forces take over. Our model and results focus entirely on the first, rapid phase.

We assume a simple trait architecture with additive biallelic loci of all the same effect size. This assumption entails a permutation symmetry that simplifies our mathematical analysis. While all loci are equivalent with respect to selection, they may differ due to the stochastic forces of mutation and genetic drift.

The strength of directional selection is arbitrary and can (and usually does) depend on time and on the genetic background via epistasis for fitness. We assume that the trait adapts either through new mutations or from mutation-selection-drift balance. Under these conditions, the adaptive architecture (the joint distribution of allele frequencies at loci contributing to phenotypic adaptation) is fully described by our analytical results, with the following main features.

A single composite parameter, the population background mutation rate Θ_bg_ emerges as the critical factor to determine the characteristics of the adaptive regime, from a sweep-type architecture for small Θ_bg_ ≪ 1 to highly polygenic ones with small shifts for Θ_bg_ ≫ 1, and oligogenic patterns in between. Qualitatively, the background mutation rate can be understood as a measure of *segregating redundancy* (Láruson *et al*. 2020): alternative genotypes to achieve phenotypic adaptation. From the standpoint of a single adaptive allele, it measures the extent of competition it faces from equivalent alleles at other loci that originate and/or rise in frequency in the time interval during which the focal allele is on its way to fixation. There are three important observations. First, the adaptive regime is independent of the strength of directional selection acting on the trait. The intuitive reason for this is that two effects of selection cancel: While the number of alternative alleles that establish per generation increases with selection (establishment probability *p*_est_∼*s*), faster growth shortens the time window for competing alleles to arise (∼1/*s* in the early phase of exponential growth). This result is analogous to the probability of soft selective sweeps, where the effects of selection on *p*_est_ and on the window of opportunity in which further beneficial alleles at the same locus can arise also cancel (Hermisson and Pennings 2005; Pennings and Hermisson 2006). Second, it does not matter whether adaptation occurs from SGV in mutation-selection-drift balance or only from new mutation. In particular, for a given Θ_bg_, adaptation from SGV is not “more polygenic”. Indeed, the heterogeneity between allele frequencies of equivalent loci in the SGV is (maybe surprisingly) large and exactly the same as the one that results from recurrent new mutation. Third, the size *L* of the genetic basis of the trait has only an indirect effect on the type of adaptation, via Θ_bg_. In particular, adaptation for a highly polygenic trait (large *L*, as would be observed in GWAS) can still be oligogenic or even sweep-like if beneficial mutation rates are low. In this case, the adaptive process can readily be described by a low-dimensional effective model with small *L* and appropriately matched Θ_bg_ (see Results).

### Adaptation dynamics in pheno-time

Our analytical method relies on a change of scale, on which the adaptive process is described. Instead of the usual time dynamics, we assess the adaptive architecture of a trait for fixed thresholds of the corresponding trait mean, 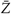, *i*.*e*., we follow “pheno-time” instead of real time. From an empirical perspective, pheno-time is a natural choice because adaptive progress at the phenotypic level is often observable. Mathematically, the change of scale reveals a robust pattern that is not visible for replicate samples that are taken after a fixed time. Due to epistasis (the selection strength changes with the distance to the optimum) and genetic drift (replicates reach phenotype thresholds at different times), both scales are not related by a simple transformation.

We observe a gradual build-up of adaptive architecture with the following steps. Any adaptive process starts with small frequency changes of alleles in the SGV: on average across replicates, the frequencies of beneficial variants slightly increase, while those of deleterious variants decrease. The adaptive architecture in this initial phase is simply a perturbed version of the joint allele frequency distribution in the balance of mutation, stabilizing selection, and drift. The phenotypic distance that is traversed in this initial phase depends on the amount of SGV and thus not only on mutation rates, but also on the strength of stabilizing selection. It can be large in the highly polygenic regime (large mutation/selection ratio, Θ_bg_/(*σγ*_2_), in our model).

Subsequently, as frequency changes get larger, deleterious variants are gradually purged from the population. The nontrivial polymorphic part of the adaptive architecture then consists purely of beneficial variants, both from SGV and from new mutation. Henceforth, the shape of the architecture is governed solely by the background mutation rate and is independent of the selection strength and the proportion of alleles that originate from SGV (which could be zero). In pheno-time, its shape follows the beneficial alleles through frequency space and converges to a quasi-stable limit once these reach fixation. Alleles that exit the frequency space by fixation are replaced by alleles at other loci that enter by new mutation, as long as there is still a sufficient supply.

The quasi-stable limit shape reflects the type of phenotypic adaptation: isolated peaks in the size-ordered marginal distributions represent successive sweeps, whereas broad distributions at many loci with strong overlaps indicate highly collective modes of adaptation. Limit architectures are quickly reached for small background mutation rates, but require adaptation across large phenotype distances in the highly polygenic case.

Finally, once adaptation progresses over distances on the order of the total phenotype range, the shape slowly changes in the direction of a less polymorphic type (smaller Θ_bg_) due to the dwindling supply of new mutations. For a trait with finitely many loci, it is therefore only quasi-stable.

All these results assume that selection is directional, and the mean trait value has not yet reached the new optimum (the *rapid phase* in Hayward and Sella 2022) Once the new adaptive optimum is reached, selection at the locus level is no longer directional, but disruptive and generally much weaker. The slow process of allele sorting that follows, and the eventual return to equilibrium at the new optimum, are not captured by our formalism.

### Theory of oligogenic adaptation

The adaptive architectures described by our framework interpolate between two regimes that provide a simplified description of the adaptive process. In the “monogenic” limit, adaptation can be conceptualized as an adaptive walk. Stochastic effects are important in this regime and lead to a heterogeneous response (complete sweeps at some loci, no response at others), but the individual steps of the walk are largely independent and can be described by single-locus theory (*e*.*g*., Kopp and Hermisson 2009b). At the other, polygenic, end of the scale, phenotypic adaptation is a collective response of alleles at very many loci. Genetic drift can be ignored, and deterministic theory can be used to describe the initial, rapid adaptive phase (*e*.*g*., Lande 1983). Loci may interact, but as long as the individual allele frequency shifts are tiny, epistatic interactions hardly play a role. In the intermediate, oligogenic regime, phenotypic adaptation is achieved by simultaneous allele frequency shifts at several, but not very many loci. In contrast to the monogenic case, adaptation is collective, and polymorphic alleles at different loci interact due to fitness epistasis. Sweeps are often partial, rather than completed, as beneficial alleles only rise to intermediate frequencies. In contrast to the highly polygenic scenario, however, frequency changes are more than just a perturbation of the standing variation. Stochastic effects are important and lead to heterogeneous contributions of otherwise identical loci (major-minor locus structure due to a mix of larger and smaller shifts).

Mathematically, oligogenic adaptation is the most challenging regime. Previous models mostly rely on deterministic theory and/or simulation studies. An insightful analytical approach is the model by Jain and Stephan (2015, 2017b,a). Building on work by de Vladar and Barton (2014), these authors study an additive QT and derive frequency trajectories of single alleles with arbitrary effects when adaptation occurs from deterministic mutation-selection balance. This initial condition favors small-effect alleles, which start from a high frequency of *p* = 0.5 (de Vladar and Barton 2014). As a consequence, the prevalence of large, sweep-like allele-frequency changes depends on the effect size, with a higher prevalence of sweeps if locus effects are large. These results extend to finite populations if and only if the population-scaled mutation rate per locus is large, Θ = 2*N*_e_ *μ >* 1, such that the distribution of weakly selected small-effect alleles in the SGV is unimodal (Devi and Jain 2023).

In our model, with low Θ and/or stronger selection, alleles typically start from a low frequency. In this case, the type of adaptive architecture (sweeps or shifts) does not depend on the effect size. Instead, the background mutation rate Θ_bg_ (which does not exist in a deterministic model) emerges as the decisive parameter. Note that our notion of a “sweep-type architecture” only refers to the size of the change in allele frequency and not to the speed: Slow frequency changes for weak-effect alleles hardly produce a discernible footprint in linked neutral variation (Thornton 2019).

The closest correspondence to our study is the analysis of a binary trait in Höllinger *et al*. (2019), where the Yule-framework has first been used. In their model, adaptation to the new optimum is achieved by a single mutational step at one of several loci underlying the trait. Many results, such as the key role of the background mutation rate Θ_bg_ and the independence of the adaptive architecture from selection strength, are equivalent in both models, demonstrating their generality. However, phenomena of multistep adaptation, such as a stable limit architecture or the effect of deleterious variation, can only be studied for a QT.

A related method, combining a Galton-Watson branching process with a deterministic logistic growth model, has recently been presented by Götsch and Bürger (2023) to study the adaptation of an additive QT under exponential directional selection from new mutation. Due to the absence of epistasis, the method can be set up in real time and allows for arbitrary locus effect sizes. Götsch and Bürger (2023) use the number of segregating alleles under selection to characterize the adaptive architecture. Once again, the total population-scaled rate of new mutations Θ_t_ (the infinite-loci counterpart of our parameter Θ_bg_) emerges as the main determinant of the pattern of adaptation.

For highly polygenic traits, further analytical approaches become available. They describe the dynamics of individual loci in a “mean field” quantitative background, which evolves according to simple deterministic dynamics. The classical approach dates back to Lande (1983) and has been used by Chevin and Hospital (2008); Chevin (2019) to describe adaptation of a major-effect allele (“QTL”) in a highly polygenic background approximated by a normal distribution. With stabilizing selection, single loci are quickly outcompeted by the joint action of the background, preventing sweep-like changes. Sweeps only occur if the shift in trait optimum is large and large-effect alleles already contribute significantly to the initial SGV (Chevin andHospital 2008; John and Stephan 2020; Stephan and John 2020; Devi and Jain 2023).

An elegant alternative approach was recently developed by Hayward and Sella (2022). Their method uses that, for an additive trait and assuming LE, selection at single loci depends on the genetic background only via the trait mean 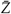 (*t*) (compare Eq. 3). For a highly polygenic trait, 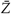 (*t*) becomes approximately independent of the dynamics at single loci. One can then insert the deterministic solution for 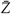 (*t*) into a single-locus diffusion equation to analyze the stochastic dynamics of individual alleles in the genetic basis. A condition for Hayward and Sella (2022)’s method is that the shift in the trait optimum is small enough that “adaptation to the new optimum requires only a small average frequency change per segregating site”. Under this assumption, all frequency changes during the rapid adaptive phase are first-order perturbations of SGV frequencies. In our model and the examples shown, these conditions are met only in the highly polygenic case of Θ_bg_ = 100 and phenotypic adaptation of no more than a few (≲ 10) steps. In this case, adaptation to the new optimum is already complete with the first step of the adaptive architecture construction described above, before its characteristic features begin to show. Accordingly, the analysis by Hayward and Sella (2022) mainly describes the relative frequency change of alleles with different effect size and the consequences for allele sorting during the equilibration phase – issues that are beyond the scope of our work.

### Oligogenic and polygenic adaptation

The transition between oligogenic and polygenic adaptation is gradual, with no clear demarcation. Besides the background mutation rate, Θ_bg_ (which determines the number of co-segregating alleles), it also depends on the range of phenotypic adaptation. Indeed, if (say) 20 beneficial alleles from the SGV are picked up by selection, adaptation can appear as “polygenic”, with small frequency shifts when the phenotype only changes by a single mutational step, but no longer at larger distances. At, say, 10 steps or more, frequency changes become large, interactions matter, and new mutations start to play a role.

A hallmark of the highly polygenic regime is that the genetic variance remains approximately constant, which is often used as a model assumption in quantitative genetic approaches (*e*.*g*., Lande 1983; de Vladar and Barton 2014; Chevin and Hospital 2008). For the infinitesimal model, a stable variance is a consequence of the even stronger assumption that selection does not change allele frequencies. As Hayward and Sella (2022) point out, it still holds for small, but non-zero frequency shifts, as long as the increase in variance due to the increase of aligned alleles (the beneficial variation) is offset by the decrease in variance due to the frequency decrease of opposing alleles (deleterious variation).

We can contrast this with characteristics of oligogenic adaptation. With a limited number of polymorphic alleles, allele frequency changes necessarily become larger. As a result, the effect of beneficial variation for changing genetic variance, as well as its contribution to the adaptive progress of the trait mean, outweighs the effect of the deleterious variation that is eventually eliminated from the population. Genetic variance typically increases during build-up of the adaptive architecture until the quasi-stable limit shape is reached. In this later phase, the genetic variance is once again approximately stable, either constant or oscillating. However, this is for different reasons than for highly polygenic adaptation over short distances: While the deleterious variation no longer plays a role, beneficial variants reach much higher frequencies. Depending on whether this frequency is smaller or larger than 0.5, further frequency increases can increase or decrease the genetic variance. In the limit, the contributions of different loci approximately cancel. The loss of variation due to fixation of alleles is compensated by recurrent new mutation, as previously described by Hill (1982a,b) for adaptation of a quantitative trait under long-term truncation selection. A comprehensive discussion of both types of stable variance can be found in Götsch and Bürger (2023).

### Biology of oligogenic adaptation

Throughout the history of quantitative genetics, evidence for highly polygenic adaptation has been gathered from various sources. This includes the classic observation that the response to artificial or natural selection on QTs is typically rapid and without major changes in genetic variance (reviewed in Sella and Barton 2019; Flatt 2020). Modern GWAS results largely supports the view of phenotypic adaptation proceeding via frequency changes at many loci with small individual effect (Shi *et al*. 2016). The contribution of genomic regions to the heritability is often roughly proportional to their length, which is consistent with an infinitesimal or “omnigenic” model (Boyle *et al*. 2017; Liu *et al*. 2019).

However, empirical evidence that goes beyond the infinitesimal model is equally widespread. In particular, major allele frequency changes driven by positive selection are frequently observed. This includes signals of sweeps and partial sweeps, but also polygenic footprints in quintessential QTs, such as human body size (Field *et al*. 2016). Such footprints do not exist with truly infinitesimal genetics.

A polygenic genetic architecture implies a high level of redundancy (Láruson *et al*. 2020; Barghi *et al*. 2020), reflecting the number of different ways how alleles can combine to produce the adaptive phenotype. As argued by Hayward and Sella (2022), it then “becomes uninteresting to focus on the particular subset of alleles” that was recruited – largely at random – to accomplish this task. However, data from replicated events of adaptive evolution often show a much higher level of parallelism than might be expected from the number of underlying loci and the segregating variation. Examples are summarized in Láruson *et al*. (2020); Barghi *et al*. (2020) and come from both natural evolution (*e*.*g*., Conte *et al*. 2015; Yeaman *et al*. 2018a) and from “Evolve and Resequence” experiments (*e*.*g*., Barghi *et al*. 2019).

Larger frequency changes and substantial parallelism are hallmarks of oligogenic adaptation: They show that the “segregating redundancy” that is available for adaptation is not unlimited. In models with equivalent loci (both for an additive and a binary trait, *cf*. Höllinger *et al*. 2019), the composite parameter Θ_bg_ = 2*N*_*e*_*μ*(*d* −1) is the appropriate measure of redundancy and determines the type of adaptation. Oligogenic characteristics are expected if (at least) one of the factors in Θ_bg_ is small. Empirically, they are highly variable: While 2*N*_*e*_*μ* measures the expected diversity per locus and strongly depends on the population size, *d* is the number of loci available to respond to a new selection pressure and depends on the size of the genetic basis. For example, melanism in flies (Bastide *et al*. 2016) or lipid traits in humans (Shi *et al*. 2016) have a much smaller genetic basis than classical size- or yield-traits.

However, it is important to distinguish the genetic basis of a trait from its “adaptive basis”, *i*.*e*., the loci that are sufficiently free of pleiotropic, epistatic, or developmental constraints to contribute to sustained adaptive change (Yeaman *et al*. 2018b; Yeaman 2022). Only the latter enter into *d* and, thus, affect Θ_bg_. Pleiotropy is necessarily ubiquitous for omnigenic traits, where trans-acting alleles in peripheral pathways with minute effects on the focal trait are thought to contribute the bulk of heritable variation (Liu *et al*. 2019). If these alleles are strongly constrained due to their primary function, the basis of the adaptive architecture could be much smaller than the total genetic basis of these traits as seen in GWAS (Láruson *et al*. 2020; Barghi *et al*. 2020).

### Scope and limits of our model

The most stringent limitation of the model concerns the assumptions on the trait genetics: alleles at all loci are additive, with equal effect sizes on the trait and on fitness. While our method describes the heterogeneity among frequencies of beneficial alleles due to mutation and drift (the stochastic variation within fitness classes), it does not capture heterogeneity due to selection differences (the deterministic variation between fitness classes). The only exception is the distinction of two classes with loci that harbor the beneficial and deleterious variation, respectively. The extension to deleterious loci shows how, in principle, multiple classes of loci can be included that differ either in their effect on the trait or in pleiotropic effects on fitness. Each additional class requires an additional model parameter and a separate fit to the starting configuration prior to the onset of directional selection. A general model with many classes can become complex, and the scope of this approach remains to be explored.

A second limitation refers to the selection regime. Our method relies on directional selection acting on all loci, as is typical during the early phase of phenotypic adaptation after a change of the optimum. There is no straight-forward extension to other modes of selection, including disruptive selection (at the locus-level) that drives adaptive fine-tuning during the equilibration phase. The assumption is also important prior to the environmental change, where standing variation is maintained by a balance of mutation and negative directional selection. For stabilizing selection, this is approximately true if selection is strong enough to keep deleterious alleles at a low starting frequency, *p*_*i*_, :S 0.1. It does not hold if much of the trait adaptation is due to common alleles that are already segregating at intermediate frequencies before adaptation, either because of very weak selection, or because they are maintained by other forces, such as balancing selection or recurrent gene flow.

Despite these restrictions, the directional selection model encompasses a large set of scenarios. In particular, our results for the adaptive architecture after a sudden shift of the trait optimum can also be applied to the second standard model of phenotypic adaptation: They hold (without change) for a trait with a moving optimum (*e*.*g*., Lynch *et al*. 1991; Bürger 2000; Jones *et al*. 2004; Kopp and Hermisson 2009a,b), as long as the gap between the optimum and the trait mean remains sufficiently large that overshooting can be ignored (the “mutation-limited regime” of Kopp and Hermisson 2009a,b). In general, the method is much less restrictive with respect to the ecological assumptions, which determine the shape of the phenotypic fitness function and its change in time, than with respect to the genetic assumptions. This includes fitness epistasis of any order (see the *Mathematical Appendix* for details), as long as the simple additive trait genetics ensures that all loci are affected in the same way.

Our analytical predictions assume LE between all loci in the genetic basis of the trait, which is unrealistic, especially for traits with a highly polygenic basis. Comparison with individual-based simulations show, however, that the results for the joint allele-frequency distribution remain accurate even for strong linkage and long-distance adaptation. This is true even though we observe (as expected, Hill and Robertson 1966) a significant increase in negative LD when the average recombination rate between pairs of loci is smaller than the selection coefficient. Although the model assumption of LE is violated, this has surprisingly little effect on the distribution of allele frequencies. Only if linkage is complete do larger deviations in the direction of a more polygenic adaptive architecture occur. It would be interesting to relate our results to the predictions from models of clonal interference (Gerrish and Lenski 1998; Desai and Fisher 2007) in this limit. Relevant effects are also expected if selection effectively acts on linkage blocks, either because initial levels of LD are large (*e*.*g*., due to admixture) or when loci underlying the trait are densely distributed along the genome (*cf*. Sachdeva and Barton 2018; Sachdeva 2022)

Finally, all our results assume a single, panmictic population. If adaptation occurs in a spatially extended population, different beneficial variants may dominate in different regions and subsequently mix through gene flow. For a single adaptive step and alleles that are mutually exclusive, the resulting spatial patterns have been analyzed (Ralph and Coop 2010; Paulose *et al*. 2019) and compared with empirical data (Feder *et al*. 2019). Furthermore, a trait optimum can vary in space and time (Polechová *et al*. 2009; Polechová 2018). All of these scenarios are expected to significantly affect patterns of adaptive architecture, both in the short term and in the long term, when gene flow drives allelic turnover (Yeaman 2015). A simpler situation in which our method could be applied is adaptation from migration-selection balance in a continent-island model. Extensions are needed to deal with strong LD due to multi-locus migration and potential barriers to gene flow. For this model, Sachdeva (2022) recently found that the effects of population-wide LD in migration-selection equilibrium can be accounted for by an adequately defined effective migration rate, providing a starting-point for a non-equilibrium analysis.

## Data availability

The associated *Dryad* directory is accessible under the DOI 10.5061/dryad.573n5tbc9 (Höllinger *et al*. 2023). This repository contains comprehensive *Mathematica* Notebooks (*Computational* and *Figures Appendix*), simulation scripts (*C++*) and created (simulation) data used for plotting. All data and scripts to recreate the presented figures are available there.

## Acknowledgments

The computational results presented have been achieved in part using the Vienna Scientific Cluster (VSC).

## Appendix

In this appendix, we present complementary figures to the figures in the main text with different choices of the parameter values. We also explain how our model extends to diploid genetics and present results for various levels of linkage in this case.

### Adaptation with LE

#### Adaptive architecture for 100 loci

Fig. S1 complements main text Fig. 4 (for a 10-locus trait) and shows the build-up of adaptive architecture for long-range adaptation of a trait with *L* = 100. Approximate convergence to a stable limit architecture can be observed for Θ_bg_ ≤10. For increasing distance of phenotypic adaptation, the results also show the broadening of the architecture (larger distances between successive size-ordered marginal distributions). As explained in the main text, this is due to the dwindling of adaptive material, corresponding to a reduction in the effective Θ_bg,eff_. The fit between LE simulation results and analytical curves is even better than in the main text Fig. 4. This is because the ≈10-fold smaller mutation rates *per locus* for *L* = 100 lead to a reduced effect of back mutation, which is ignored in the approximation.

#### Dynamics of summary statistics

Adding to Fig. 5 for a 10 000-locus trait in the main text, we show summary statistics of the adaptive architecture of QTs with 10 and 100 loci in Fig. S2. The adaptive scenarios correspond to the ones of Fig. 4 and Fig. S1, respectively. In addition, we also show a case of adaptation from only new mutation (*de novo*) for *L* = 10. Furthermore, we show how our results for pheno-time relate to the usual dynamics in real time.

The first row of Fig. S2 shows the scaled number of generations (on the *y*-axis, in units of Θ_bg_ *σγ*^2^) required for the mean phenotype 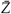 to reach adaptive thresholds in units of mutational steps *c*_Z_ (*x*-axis). To derive a simple analytical prediction for this dependence, we use a result by Götsch and Bürger (2023) for an additive trait under constant directional selection and constant rates of beneficial mutation (assumption of an infinite-locus model). Translated to our model parameters, Götsch and Bürger (2023, Eq. 4.21) derive an expected change in the mean trait of

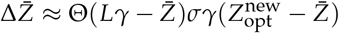

for long-term adaptation. That is, to leading order in the selection coefficient, the average adaptive progress in the trait mean is simply given by the product of the selection strength and the population mutation rate. In our model, both the selection strength (which scales with the distance to the new optimum, 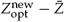) and the rate of new beneficial mutations (which is proportional to the distance to the end of the phenotype range, 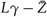) are variable. This leads to a differential equation, which, in dimensionless quantities, can be written as

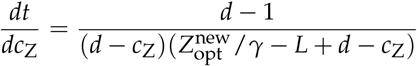

where time *t* is measured in units of [Θ_bg_ *σγ*^2^]^−1^. The solution is

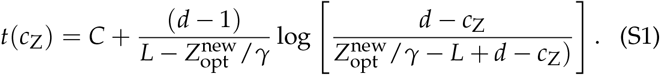

This curve is displayed as a black dashed line in the first row of Fig. S2 (with *C* chosen such that it passes through (0, 0)). Similarly, the analytical expectations (dashed curves) for the equilibrium variance in the second row of Fig. S2 are based on Götsch and Bürger (2023, Eq. 4.24). In terms of our model parameters and with a variable mutation rate, this reads, *d* − 1

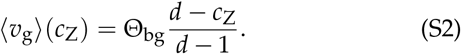

We observe the following:

The analytical prediction (Eq. S1) (which assumes an infinite-locus model) matches the simulation data in the top row whenever the background mutation rate Θ_bg_ is small enough that recurrent mutation at the same locus is unlikely. For small *L* and/or high Θ_bg_, new mutations frequently occur at loci where mutant alleles are already established, which results in slower dynamics (steeper increase).

For the variance (second row), we see that the estimate for the average variance in real time (Eq. S2) is consistently lower than the simulation data for small Θ_bg_ *<* 1. The reason (also observed by Götsch and Bürger 2023) is that sampling at fixed phenotype thresholds does not represent time averages. Indeed, for very small Θ_bg_, there is no segregating variation for the trait most of the time, which is only possible for sampling at integer values of *c*_Z_ (where the approximation matches more closely). For higher Θ_bg_, the pheno-time values no longer oscillate and approach the time average – until saturation effects due to recurrent mutation at the same loci lead to lower variances in a finite-locus model compared to an infinite-locus model for large Θ_bg_. See also Fig. 5 for *L* = 10 000 in the main text, where the real time averages from the infinite-locus model (*v*_g_≈ Θ_bg_) are approximately met once the oscillations become small for Θ_bg_ ≥1.

The initial variance in SGV is smaller than the equilibrium long-term variance in all cases shown, even for Θ_bg_ = 100. Until the dynamic equilibrium is reached, the rate of adaptation is slower (which leads to a shift to larger *y*-values in the top row, see Θ_bg_ = 10 for *L* = 100 and Θ_bg_ = 1 for *L* = 10). Since initial variation is absent in the *de-novo* scenario, the rate of adaptation is even slower (compare first and second column, top row). However, differences in variance and skew due to the starting conditions vanish quickly (≲ 1 mutational step for Θ_bg_ ≤10).

Positive initial skew (bottom row) results from the asymmetric starting conditions (more mutations to increase the trait than to decrease it). Negative genetic skew indicates a dwindling supply of fresh loci, which is either transient (yielding oscillations) for low effective Θ_bg,eff_, or persistent when the finite trait basis has been exhausted.

### Adaptation with linkage

#### Haploids with linkage

We extend Fig. 6 to other settings of Θ_bg_: 0.01 (Fig. S3), 0.1 (Fig. S4), 10 (Fig. S5) and 100 (Fig. S6). The effect of linkage is strongest for intermediate values of Θ_bg_:

**Figure S1.**
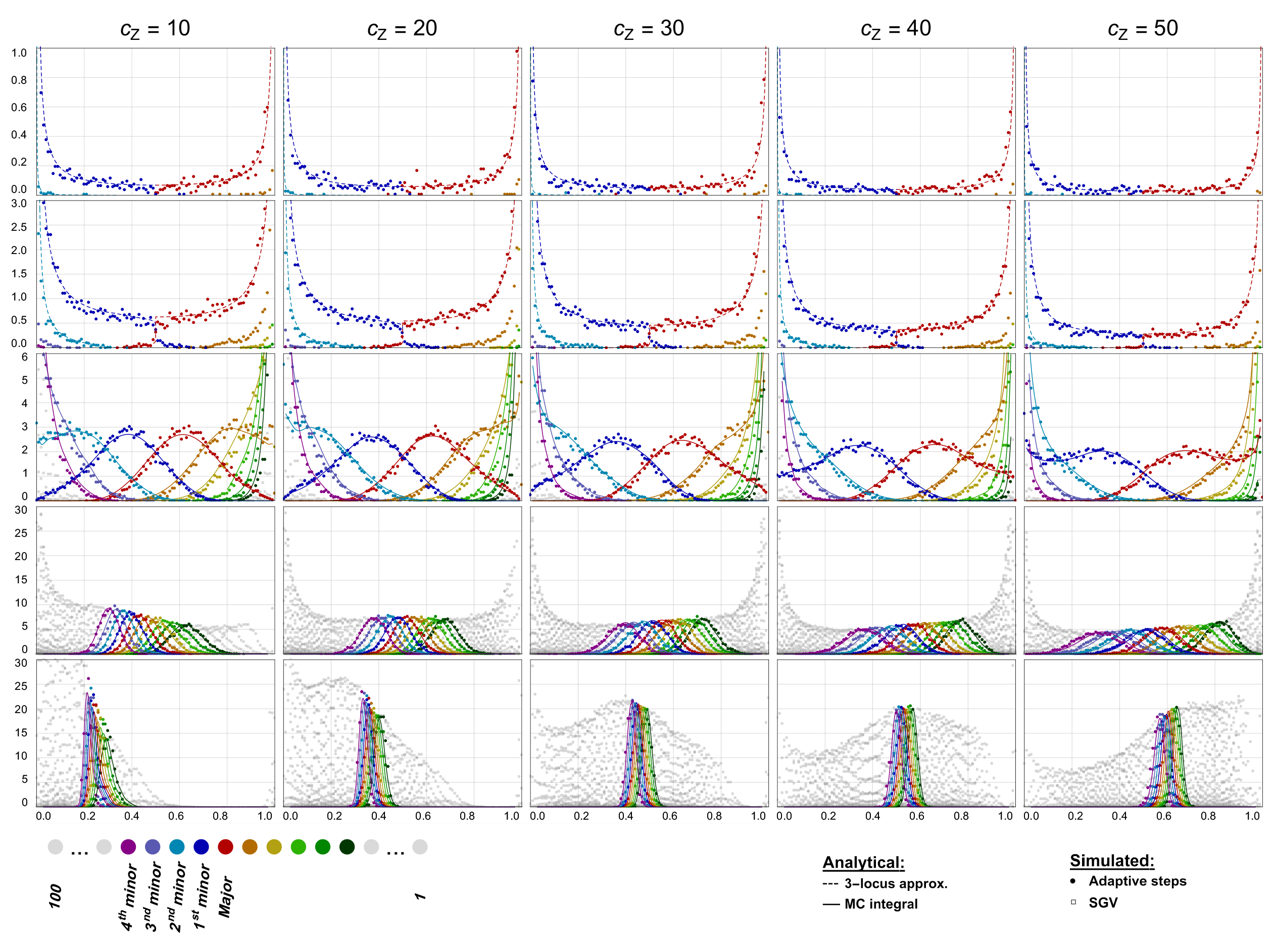
Dynamics of the adaptive process for a QT with 100 loci. Successive snapshots (in intervals of 10 mutational steps) of the ordered marginal allele-frequency distributions over the course of adaptation of a QT with 100 loci. Equilibrated populations evolve from an initial optimum 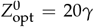 toward a new optimum 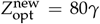. LE simulation densities (points) are accurately predicted by the analytical model. Dashed curves (for Θ_bg_ = 0.01 and 0.1) show analytical three-locus approximations accounting for the progressive reduction in the effective Θ_bg,eff_ with *c*_Z_ (see main text). Solid lines correspond to results of the 100-locus Monte-Carlo method, including deleterious variation (*cf*. the *Mathematical Appendix*). The distributions of the loci with the 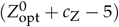th (dark green) to 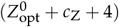th (magenta) largest frequency are shown in bright colors. All others are shown in gray. We include a step-change in the strength of stabilizing selection from *N*_e_ *σ*(*t*)*γ*^2^ = 100 for *t <* 0 to 10 for *t* ≥ 0 to avoid artifacts due to excessively strong directional selection for *t* ≥ 0. *N*_e_ = 10 000, 10 000 replicates. Note the differences in scaling of the y-axes.

**Figure S2.**
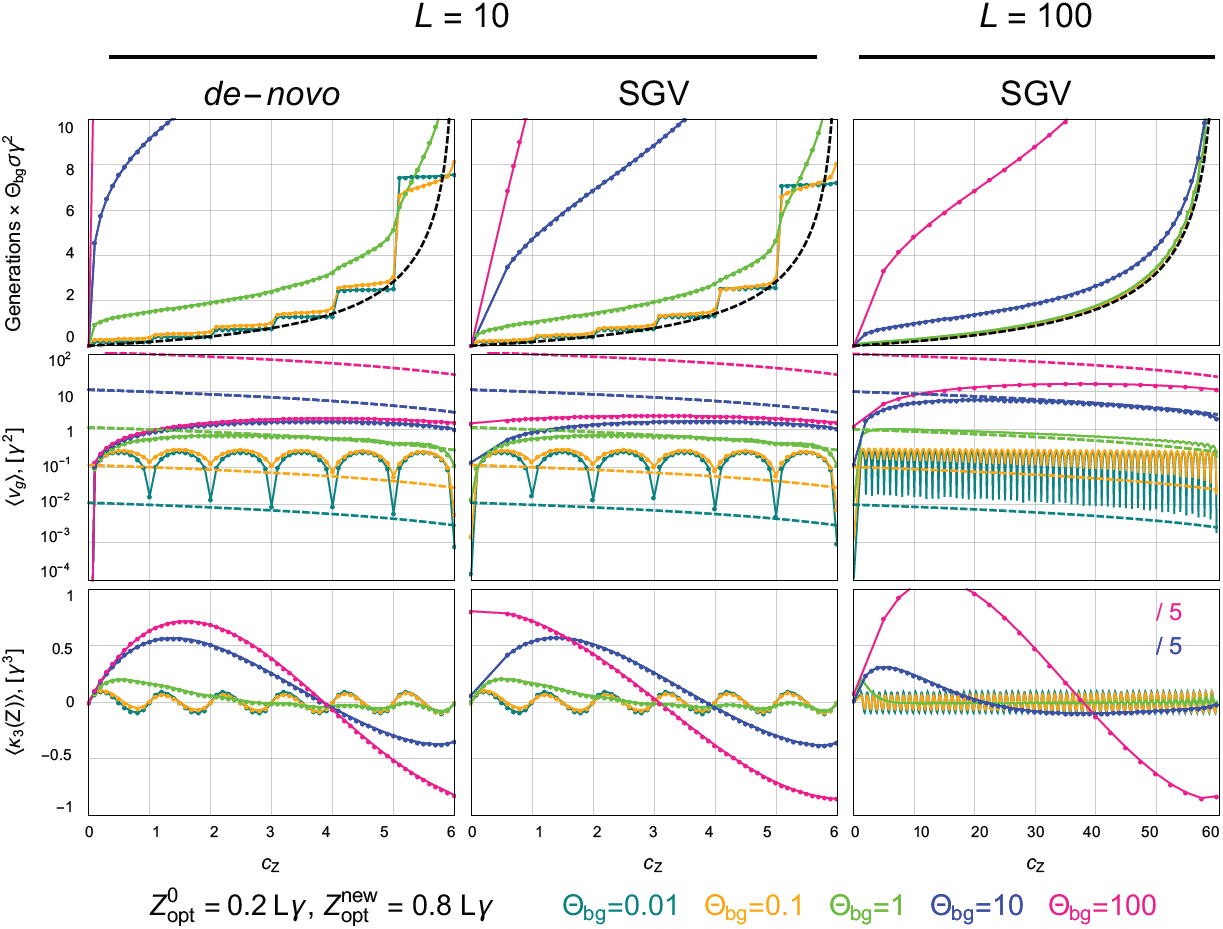
Genetic variance and skew for QTs with 10 and 100 loci. QTs with 10 loci (two left columns: without/with SGV) and 100 loci (right: with SGV) adapt from 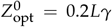 to 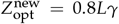 (6 mutational steps for *L* = 10 and 60 steps for *L* = 100). The top row shows the average number of generations to reach thresholds of phenotypic adaptation (real time in terms of pheno-time). The dashed black line indicates an approximate theoretical expectation (for all Θ_bg_, see Eq. S1). The middle and bottom row show the dynamics of the genetic variance (with expectations as dashed lines, see Eq. S2) and the skew in pheno-time. *N*_e_ = 10 000, *N*_e_ *σγ*^2^ = 100 (switch to *N*_e_ *σγ*^2^ = 10 for *t >* 0 for 100 loci, *cf*. Fig. S1), 10 000 replicates (1 000 and 5 000 for Θ_bg_ = 0.01 resp. 0.1 in first row). The insets “/5” indicate that the corresponding lines are 1/5th of the actual data.

- For low mutation rates, tightly linked loci are rarely polymorphic simultaneously. For Θ_bg_ = 0.01 (Fig. S3), the adaptive architecture remains unaltered even for *r* = 0, while for Θ_bg_ = 0.1 (Fig. S4) slight changes are visible.
- While the effect of linkage for Θ_bg_ = 10 (Fig. S5) is similar compared to Θ_bg_ = 1 (Fig. 6), all differences vanish for Θ_bg_ = 100 (Fig. S6). With very high mutation rates, the randomizing effect of recurrent, bi-directional mutation eliminates LD.

For large Θ_bg_ ≥10, the first phenotypic checkpoint is placed at *c*_Z_ = 2 since the trait mean in the SGV exceeds *c*_Z_ = 1 in some replicates (with asymmetric starting condition, mutation biases the trait mean upwards).

#### Diploids with linkage

For simplicity, we have derived and presented all our results so far for a haploid population. Here we show how the formalism extends to diploids without dominance. To match the haploid dynamics, we consider a diploid model with the same number of gametes, 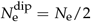 and maintain the same mutation rates *μ* per locus, such that 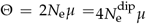. We assume the same number of loci *L* underlying the trait and the same proportion of loci with *a*_*i*_ -majority vs. *A*_*i*_ - majority allele at the initial optimum 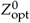 (constant *d*). We thus have 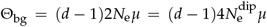. To maintain the same level of SGV (with expected frequency 2*μ*/(*σγ*^2^)), we also hold the selection strength *σγ*^2^/2 prior to the environmental change constant (compare the first columns, “SGV”, between Fig. 6 and Fig. S7).

Lastly, we also want to make the genic, 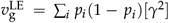, and genetic variance, *v*_g_ = ∑_*Z*_ *Z*^2^ *q*_*Z*_ − (∑_*Z*_ *Zq*_*Z*_)^2^ (where *q*_*Z*_ is the frequency of phenotype *Z* in the population), comparable between haploids and diploids (see insets in Figs. 6 and S7). To that end, we need to set 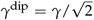 because the trait is a sum over the contributions of 2*L* = 20 alleles in the diploid case, instead of *L* = 10 for haploids. Accordingly, the phenotype range is 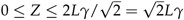 and the optimum shifts by 6 mutational steps of size 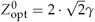 from 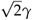 to 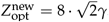.

**Table S1.** Nomenclature. Overview of the main parameters used in the manuscript.

| Symbol | Meaning | Formula |
| --- | --- | --- |
| $N_e$ | (effective) population size | |
| $p_i$ | frequency of allele $A_i$ at locus $i$ | $0 \leq p_i \leq 1$ |
| $\gamma, \mu_i$ | effect of allele $A_i$ (all equal) and locus mutation rate (usually equal, $\mu_i = \mu$) | |
| $L, d$ | total number $L$ of loci in the genetic basis, $d$ of which have $p_i(0) \leq 1/2$ | |
| $Z, \bar{Z}$ | trait value and its mean<sup>a</sup> | $\bar{Z} = \sum_{i=1}^L \gamma p_i$ |
| $v_g, \kappa_3(Z)$ | genetic variance and skew of $Z$ | |
| $Z_{\text{opt}}^0, Z_{\text{opt}}^{\text{new}}$ | initial trait optimum ($t < 0$), new trait optimum ($t \geq 0$) | $Z_{\text{opt}}^0 = d\gamma$ |
| $c_Z$ | stopping condition (sampling point) for $\bar{Z}$<sup>b</sup> | $\bar{Z} = Z_{\text{opt}}^0 + c_Z \gamma \leq Z_{\text{opt}}^{\text{new}}$ |
| $\Theta_{\text{bg}}$ | population-scaled background mutation rate ($\mu = \mu_i$) | $\Theta_{\text{bg}} = 2N_e\mu(d-1)$ |
| $W(Z), w(Z)$ | Wrightian (discrete-time) and Malthusian (continuous-time, $\ln W(Z)$) fitness | $W(Z) = \exp \left[ -\frac{\sigma}{2} (Z - Z_{\text{opt}})^2 \right]$ |
| $\sigma\gamma^2$ | selection coefficient against minority-allele in SGV | $s_{\text{eq}} = \sigma\gamma^2 = s(t), t < 0$ |

**Figure S3.**
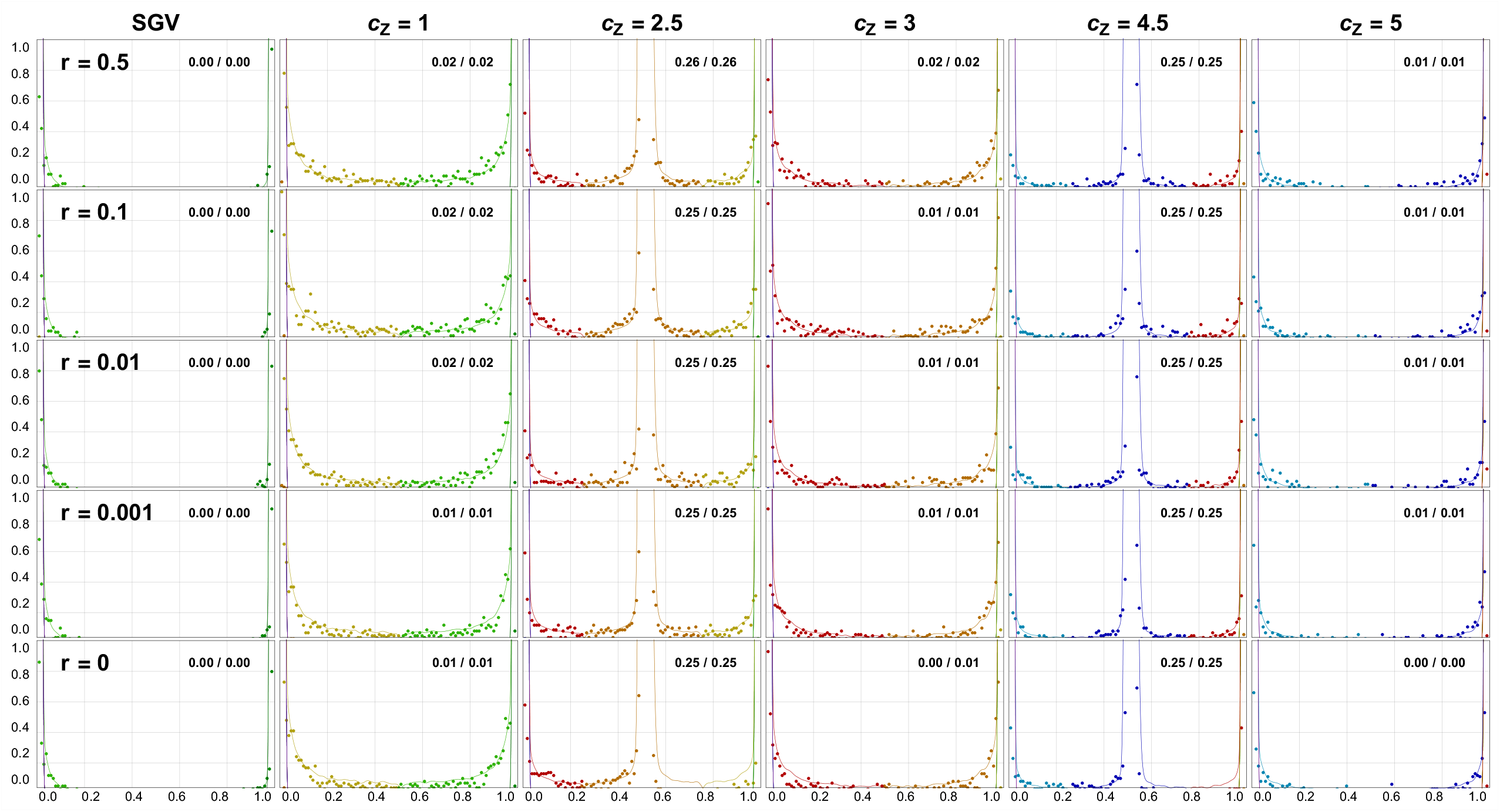
Effect of linkage on adaptive architecture of a QT (haploids, Θ_bg_ = 0.01). With very low background mutation rates, only complete linkage causes apparent changes in the adaptive architectures. Other parameters are as in Fig. 6.

**Figure S4.**
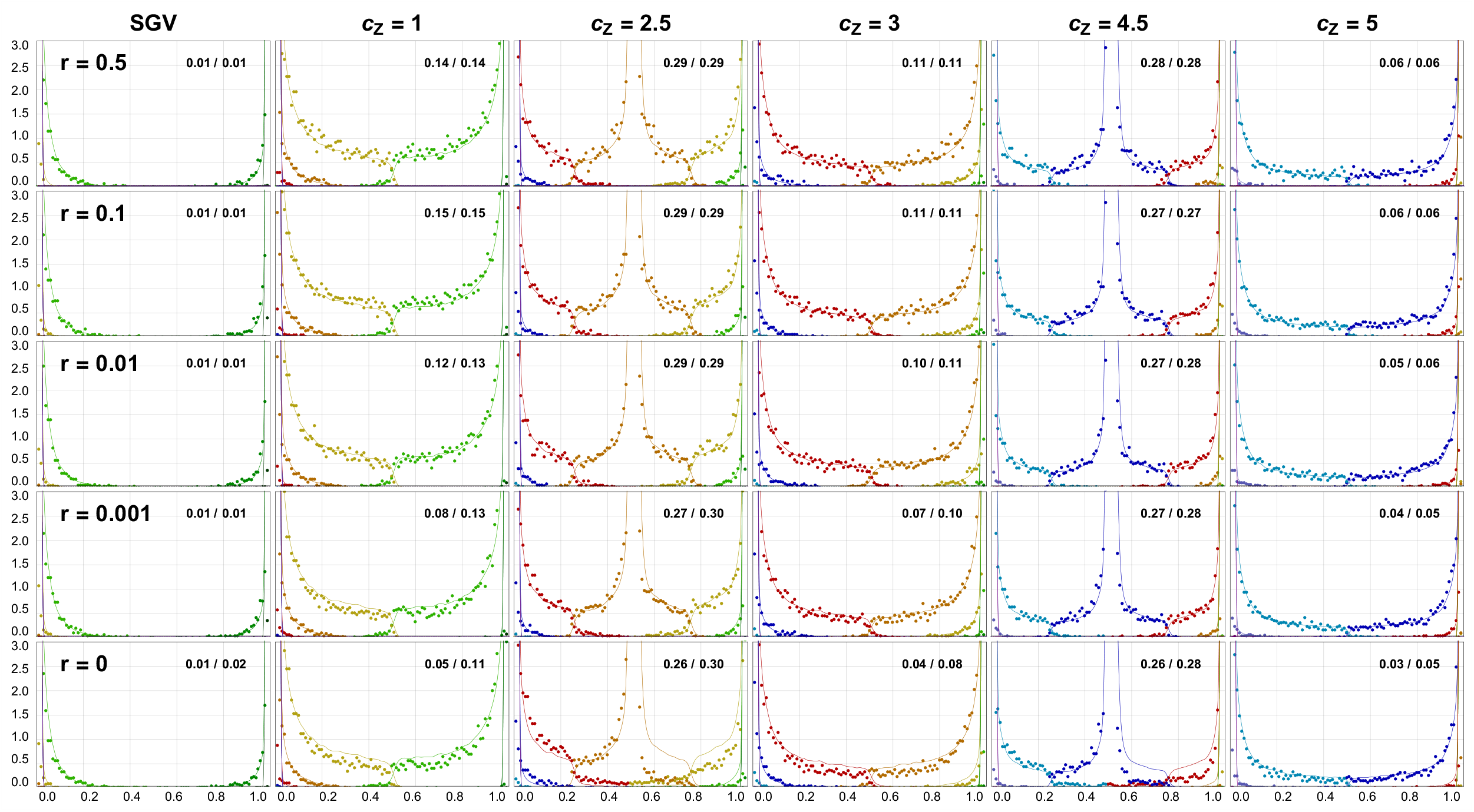
Effect of linkage on adaptive architecture of a QT (haploids, Θ_bg_ = 0.1). With low background mutation rates, the adaptive architectures are still highly robust, even under tight linkage. Other parameters are as in Fig. 6.

**Figure S5.**
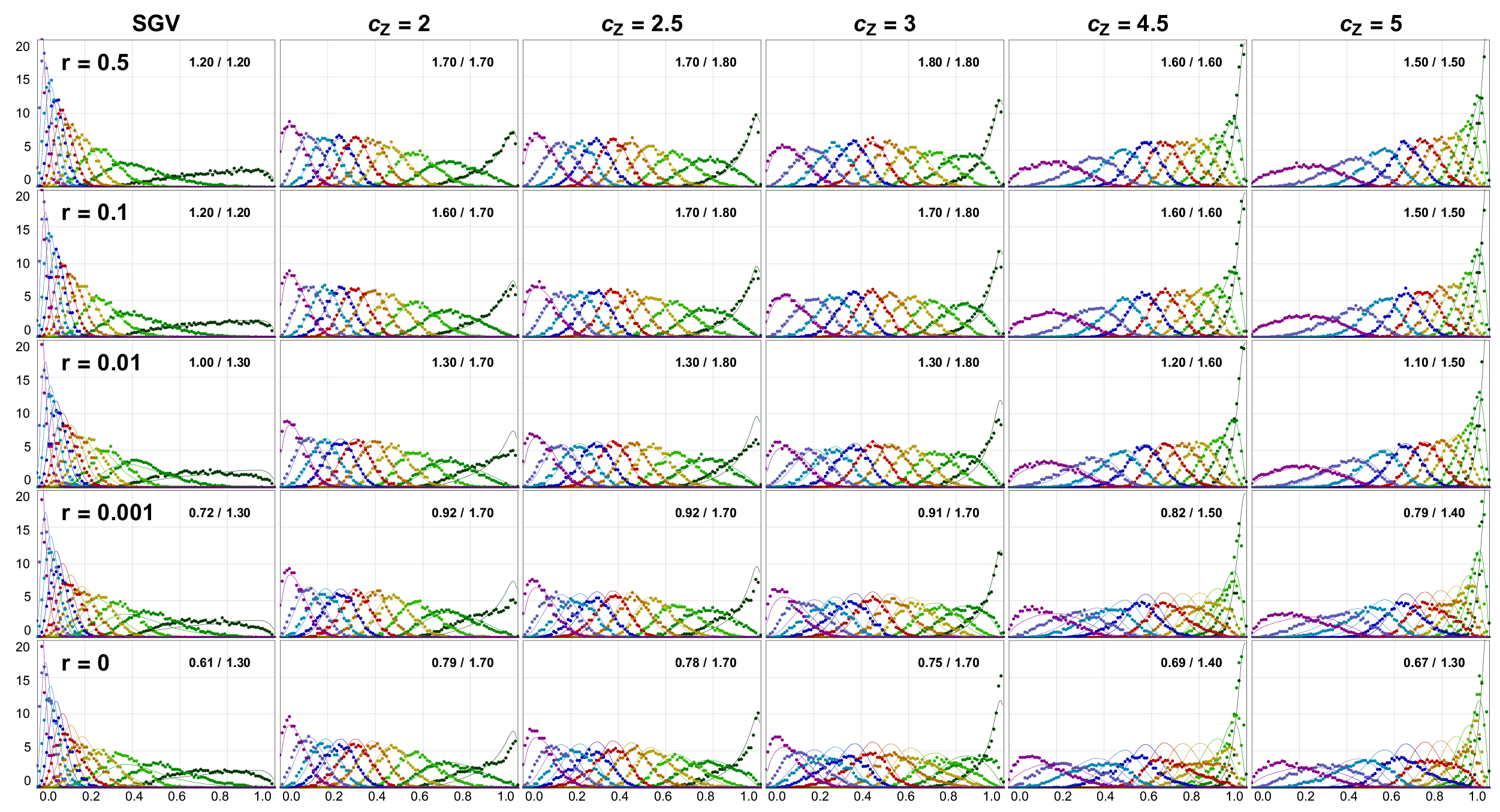
Effect of linkage on adaptive architecture of a QT (haploids, Θ_bg_ = 10). With high background mutation rates, we can detect mild alterations under tight and complete linkage. Other parameters are as in Fig. 6.

**Figure S6.**
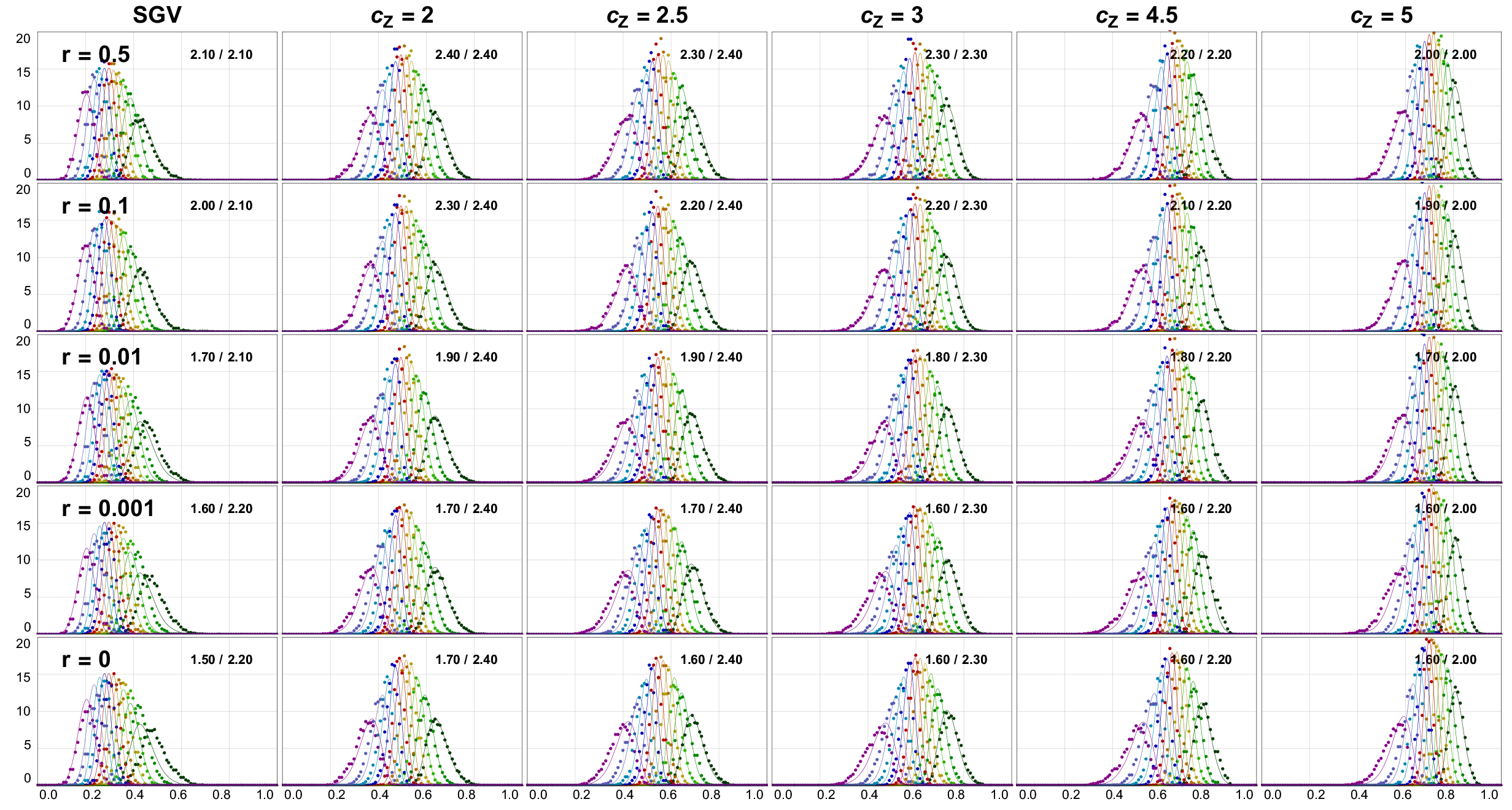
Effect of linkage on adaptive architecture of a QT (haploids, Θ_bg_ = 100). With very high background mutation rates, linkage has no apparent effect on the adaptive architectures. Other parameters are as in Fig. 6.

**Figure S7.**
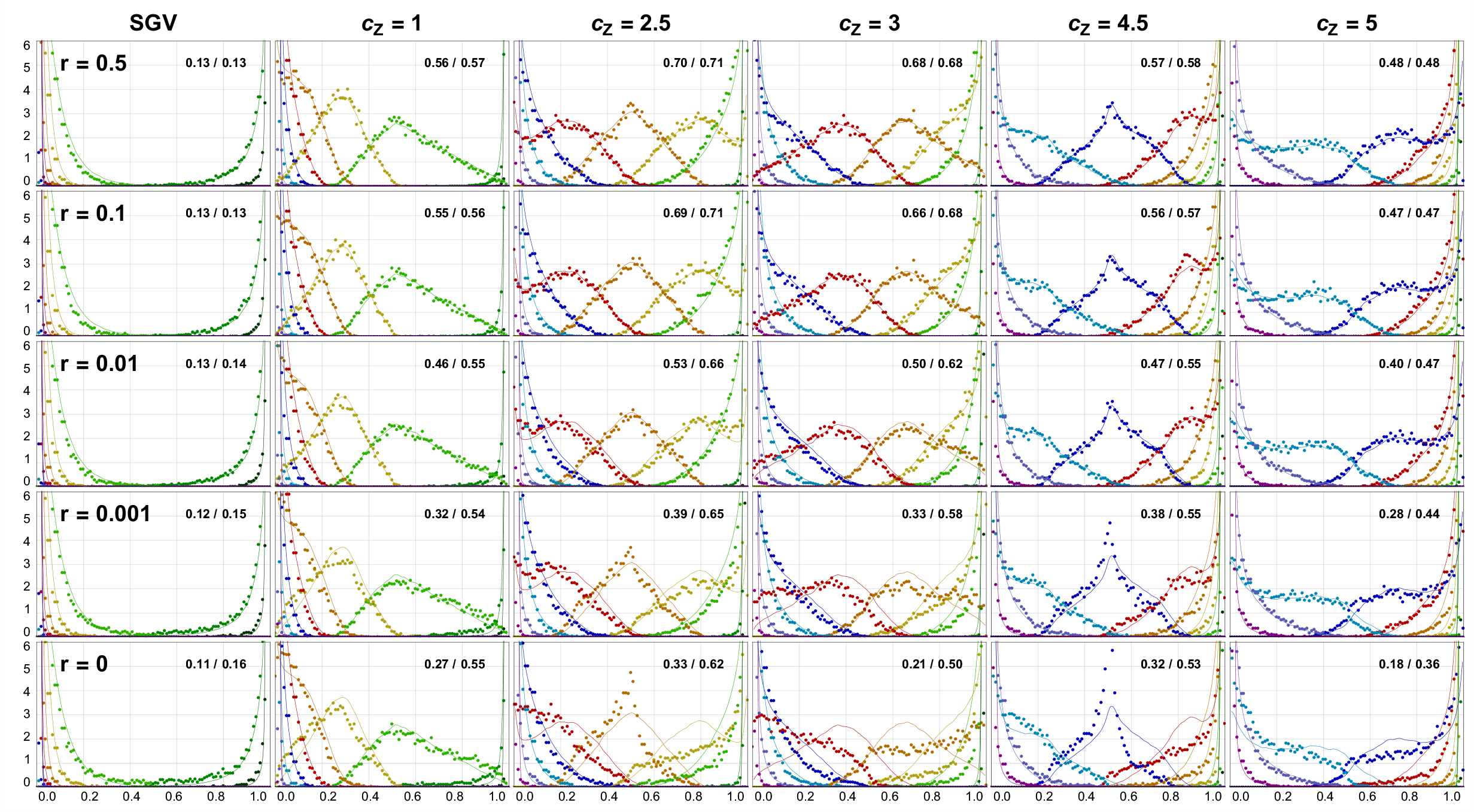
Effect of linkage on adaptive architecture of a QT (diploids, Θ_bg_ = 1). The shape of the marginal distributions is virtually identical for diploids without dominance 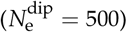 and haploids (*N*_e_ = 1000, Fig. 6). Importantly, *c* corresponds to steps of size 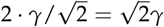. All other parameters and the curves (LE) are the same as in Fig. 6.

